# Distinct input-specific mechanisms enable presynaptic homeostatic plasticity

**DOI:** 10.1101/2024.09.10.612361

**Authors:** Chun Chien, Kaikai He, Sarah Perry, Elizabeth Tchitchkan, Yifu Han, Xiling Li, Dion Dickman

## Abstract

Synapses are endowed with the flexibility to change through experience, but must be sufficiently stable to last a lifetime. This tension is illustrated at the *Drosophila* neuromuscular junction (NMJ), where two motor inputs that differ in structural and functional properties co-innervate most muscles to coordinate locomotion. To stabilize NMJ activity, motor neurons augment neurotransmitter release following diminished postsynaptic glutamate receptor functionality, termed <u>p</u>resynaptic <u>h</u>omeostatic <u>p</u>otentiation (PHP). How these distinct inputs contribute to PHP plasticity remains enigmatic. We have used a botulinum neurotoxin to selectively silence each input and resolve their roles in PHP, demonstrating that PHP is input-specific: Chronic (genetic) PHP selectively targets the tonic MN-Ib, where active zone remodeling enhances Ca^2+^ influx to promote increased glutamate release. In contrast, acute (pharmacological) PHP selectively increases vesicle pools to potentiate phasic MN-Is. Thus, distinct homeostatic modulations in active zone nanoarchitecture, vesicle pools, and Ca^2+^ influx collaborate to enable input-specific PHP expression.

## INTRODUCTION

Synapses have the remarkable ability to adaptively adjust their strength in response to the myriad challenges they confront during development, maturation, experience, and disease. These processes, referred to collectively as “homeostatic synaptic plasticity”, have been characterized in diverse invertebrate and mammalian nervous systems(Pozo and Goda, 2010; Wang and Rich, 2018; Frank et al., 2020). Most studies have examined homeostatic plasticity in cells innervated by multiple neurons, and it has been difficult to disambiguate which specific inputs are undergoing homeostatic plasticity, their temporal characteristics, and the pre- vs post-synaptic mechanisms involved. Indeed, the vast complexity of neural circuits, where each neuron can be innervated by hundreds of other neurons forming thousands of individual synapses that differ in strength, subtype (excitatory, inhibitory, neuromodulatory), and mode (ionotropic vs metabotropic), pose major challenges towards gaining a clear understanding of how individual cells embedded within circuits are homeostatically controlled. Hence, while we have learned much about the general mechanisms mediating diverse forms of homeostatic plasticity including synaptic scaling(Turrigiano et al., 1998; Turrigiano, 2008; Li et al., 2019), firing rate plasticity(Marder, 2011; Turrigiano, 2012), and presynaptic homeostatic plasticity(Davis and Muller, 2015), how distinct inputs are selectively controlled to orchestrate circuit stability remains obscure.

In principle, the *Drosophila* larval neuromuscular junction (NMJ) is a powerful system to resolve input-specific mechanisms of homeostatic plasticity at excitatory synapses. At this model glutamatergic synapse, most muscles are co-innervated by two motor inputs, the “tonic” MN-Ib and “phasic” MN-Is, that differ in structural and functional properties(Johansen et al., 1989; Kurdyak et al., 1994; Lnenicka and Keshishian, 2000; Aponte-Santiago and Littleton, 2020). Synaptic strength at this NMJ is stabilized in response to perturbations that diminish postsynaptic glutamate receptor (GluR) functionality through a retrograde signaling system that adaptively enhances presynaptic neurotransmitter release(Delvendahl and Muller, 2019; Goel and Dickman, 2021), a process termed <u>p</u>resynaptic <u>h</u>omeostatic <u>p</u>otentiation (PHP). Two ways of inducing PHP expression have been extensively characterized: “acute PHP” refers to the pharmacological blockade of postsynaptic GluRs, which rapidly induces PHP expression within 10 mins(Frank et al., 2006), while “chronic PHP” refers to the genetic ablation of a subset of GluRs, leading to a long-term expression of PHP(Petersen et al., 1997). To date, dozens of genes have been identified to be necessary for both acute and chronic PHP expression(Frank, 2014; Goel and Dickman, 2021), which together ultimately function to enhance 1) presynaptic Ca^2+^ influx and 2) the number of synaptic vesicles available for release(Weyhersmuller et al., 2011; Muller and Davis, 2012; Goel and Dickman, 2021). Although much has been learned about the genes and mechanisms that enable acute and chronic PHP expression, how PHP signaling adaptively modulates presynaptic function at the tonic MN-Ib and/or the phasic MN-Is has not been resolved.

Recent efforts at characterizing input-specific PHP expression at tonic vs phasic synapses have led to conflicting results and interpretations. The first study to suggest that at least chronic PHP might operate with input specificity used a “quantal” Ca^2+^ imaging approach, where enhanced glutamate release was selectively observed at tonic MN-Ib synapses(Newman et al., 2017). However, a later study used selective optogenetic stimulation to conclude that both tonic and phasic motor inputs underwent PHP modulation during acute and chronic PHP, concluding that PHP non-discriminatorily targeted both inputs(Genç and Davis, 2019). While these studies made important observations, there were several limitations that rendered clear interpretations about input-specific PHP signaling difficult. First, while quantal Ca^2+^ imaging can assess input-specific differences in quantal size and transmission, this approach lacks the sensitivity and robust electrophysiological techniques necessary to understand the changes in presynaptic function that enable PHP expression. Second, the chronic expression of channel-rhodopsin necessary for selective optogenetic stimulation perturbs synaptic transmission at the fly NMJ(Han et al., 2022), does not permit some sophisticated electrophysiological assays, and is unable to resolve input-specific differences in quantal size essential to properly understand GluR perturbation and PHP expression.

Recently, a new approach was developed that enables electrophysiological isolation of neurotransmission from MN-Ib vs -Is, where selective expression of a <u>bo</u>tulinum <u>n</u>eurotoxin (BoNT-C) blocks all transmission without inducing toxicity or heterosynaptic plasticity from the convergent motor neuron(Han et al., 2022; He et al., 2023). Here, we have used selective BoNT-C silencing of MN-Ib vs -Is to determine whether acute and/or chronic PHP happens input-specifically and to interrogate the mechanisms involved. This approach has clearly demonstrated that at physiologic Ca^2+^ levels and below, PHP expression is distinctly input-specific: Chronic PHP is indeed selectively expressed at tonic MN-Ib synapses, while acute PHP is only observed at phasic MN-Is synapses. Additional electrophysiological, Ca^2+^ imaging, and confocal and super-resolution experiments demonstrate that distinct expression mechanisms are recruited to either input to enable PHP expression: Chronic PHP remodels active zones to enhance Ca^2+^ channel abundance at tonic MN-Ib release sites, leading to increased Ca^2+^ influx and neurotransmitter release. In contrast, acute PHP “compacts” active zone nanostructures to recruit more synaptic vesicles available for release. Together, PHP signaling distinctly transforms motor inputs to enable the homeostatic stabilization of muscle excitation across diverse synaptic subtypes.

## RESULTS

### Distinct motor neurons selectively express chronic and acute PHP

We first set out to unambiguously resolve whether acute and/or chronic PHP is expressed at either (or both) motor inputs at the *Drosophila* NMJ. Previously, we engineered transgenic expression of <u>bo</u>tulinum <u>n</u>eurotoxin C (BoNT-C) and demonstrated that expression of BoNT-C in motor neurons effectively silences both spontaneous (miniature) and evoked neurotransmission without confounding changes in NMJ structure or function from the convergent input(Han et al., 2022). Using selective silencing of MN-Ib or -Is by input-specific expression of BoNT-C, we went on to show that transmission from tonic MN-Ib inputs facilitate, where active zones are large, abundant, and function with relatively low release probability (P_r_) characteristics(He et al., 2023). In contrast, transmission from phasic MN-Is depresses, saturating at physiological Ca^2+^ concentrations (1.8 mM), and contains fewer active zones that are smaller in area and function with relatively high P_r_(He et al., 2023). We used this same approach as a foundation to now assess input-specific PHP expression.

We first examined chronic PHP expression at isolated MN-Ib and -Is NMJs across a range of extracellular Ca^2+^ concentrations. As previously reported, chronic PHP is observed in *GluRIIA* null mutants at ambiguated NMJs (stimulation of both Ib+Is), where stable evoked amplitudes are observed across a range of Ca^2+^ conditions (0.4 – 6 mM; **Fig. 1A,B** and Table S1). At disambiguated NMJs, weak MN-Ib NMJs contribute ∼1/3 of the evoked amplitude, while stronger MN-Is inputs confer ∼2/3 of the transmission under physiological (1.8mM) Ca^2+^ concentrations and below(Han et al., 2022; He et al., 2023) (**Fig. 1A** and Table S1). Genetic loss of the *GluRIIA* subunit leads to reduction of miniature amplitude (quantal size) by ∼50% across all Ca^2+^ levels due to loss of one of the two receptor subtypes, as expected (**Fig. 1B** and Table S1). However, at isolated MN-Ib in physiological extracellular Ca^2+^ conditions (1.8mM), presynaptic glutamate release (quantal content) is increased almost 300% in *GluRIIA* mutants compared to wild type, leading to enhanced evoked EPSC amplitude above MN-Ib baseline values (**Fig. 1A,B**). In contrast, EPSCs are reduced at isolated MN-Is NMJs by ∼50% in *GluRIIA* mutants compared to wild-type Is controls (**Fig. 1A,B**). Notably, the large 200% increase in quantal content at the weak MN-Ib compensated for reduced transmission at the strong Is to effectively stabilize overall (Ib+Is) transmission. This suggests that chronic PHP is expressed exclusively at MN-Ib, while apparently no functional changes are observed at MN-Is at physiological Ca^2+^ levels.

**Figure 1:**
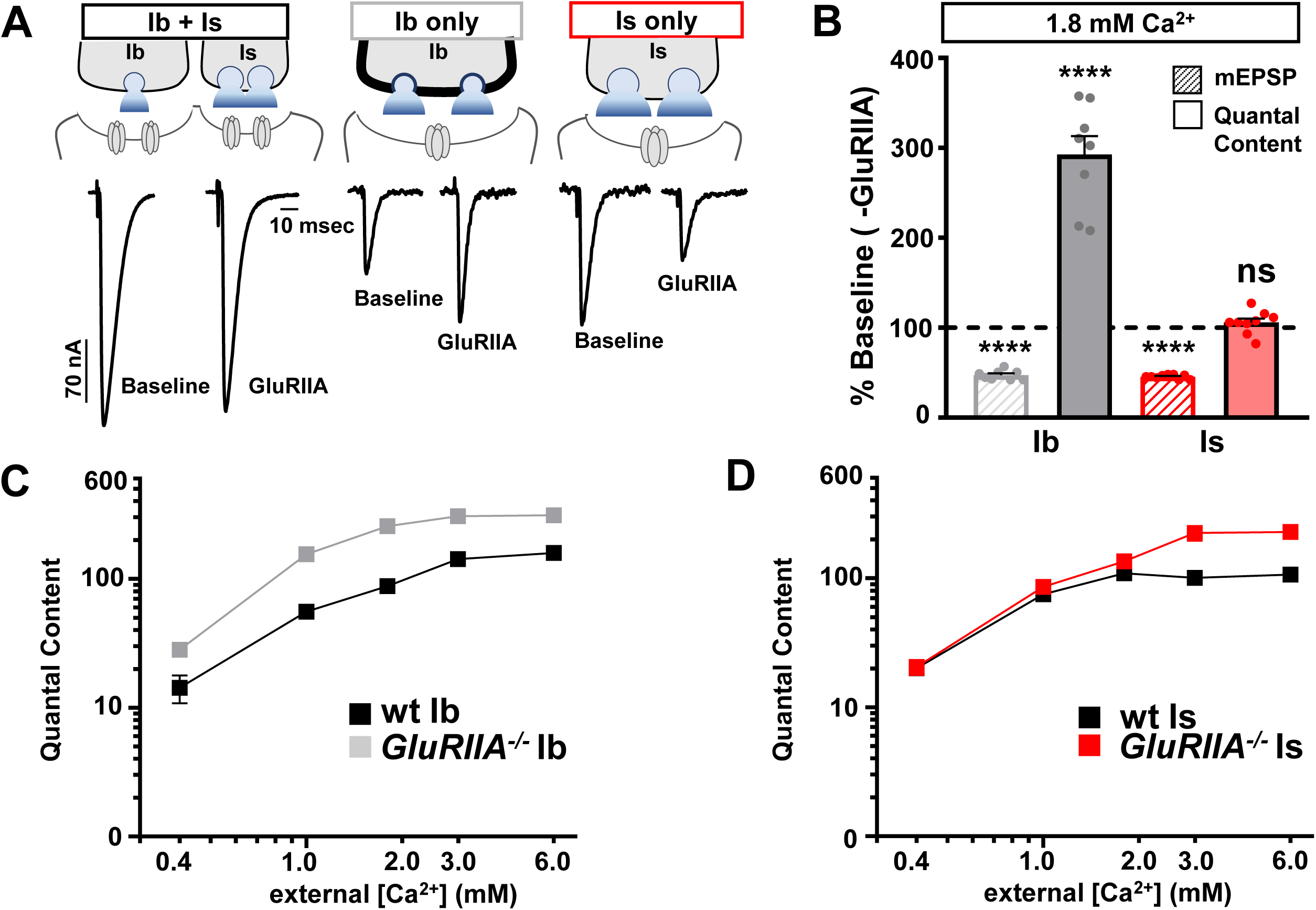
Chronic PHP is selectively expressed at tonic MN-Ib synapses. **(A)** Schematics and representative EPSC traces for baseline (wild type; *w^1118^*) and chronic PHP (GluRIIA^-/-^; *w*;*GluRIIA^pv3^*) NMJs at isolated MN-Ib and -Is inputs following selective expression of BoNT-C at 1.8 mM extracellular [Ca^2+^]. **(B)** Quantification of mEPSP amplitude and presynaptic glutamate release (quantal content) at MN-Ib and -Is in *GluRIIA* mutants normalized to wild type. Note that while quantal content is enhanced at MN-Ib, characteristic of robust PHP expression, no change is observed at MN-Is. **(C,D)** Plot of quantal content as a function of external [Ca^2+^] for MN-Ib and -Is at baseline (wild type) and after chronic PHP signaling (GluRIIA^-/-^) from MN-Ib (C) and MN-Is (D). Note that the enhanced quantal content characteristic of PHP saturates at 1.8 mM Ca^2+^ and above at MN-Ib, while quantal content does not increase at MN-Is at physiological Ca^2+^ and below. Error bars indicate ± SEM. Additional statistical details are shown in Table S1.

Differences in release probability, presumably due to changes in extracellular Ca^2+^/Mg^2+^ concentrations, were speculated to explain the differences in previous attempts to resolve input-specific PHP expression(Newman et al., 2017; Genç and Davis, 2019). We therefore performed the same experiments across a series of Ca^2+^ levels. We found that chronic PHP remained expressed exclusively at MN-Ib at physiological Ca^2+^ conditions and below (0.4, 1.2, and 1.8 mM; **Fig. 1C,D**). While quantal content does not change at MN-Is at physiological Ca^2+^ and below in *GluRIIA* mutants compared to wild type, we did note an apparent increase in quantal content at Is in *GluRIIA* mutants at high (non-physiological) Ca^2+^ levels (3 and 6 mM; **Fig. 1D**). It is difficult to interpret this result, since release from wild-type MN-Is saturates at physiological Ca^2+^ levels(He et al., 2023). Nevertheless, chronic PHP is input-specific at physiologic Ca^2+^ conditions at below, exclusively targeting MN-Ib for PHP plasticity.

Next, we performed the same set of experiments in isolated MN-Ib vs -Is NMJs at baseline and following PhTx application to block glutamate receptors and assess acute PHP expression. In these experiments, we found that acute PHP selectively targets the opposite motor input, MN-Is, for homeostatic modulation: Quantal content is enhanced after PhTx application across all Ca^2+^ concentrations at MN-Is NMJs (0.4 – 6 mM) compared to baseline MN-Is, while essentially no change is observed at MN-Ib+PhTx at physiological Ca^2+^ and below (0.4 – 1.8mM) compared to baseline values (**Fig. 2A-D**). Notably, quantal content did not need to be as robustly enhanced at the strong Is input to compensate for loss of transmission from the weak Ib to maintain stable Ib+Is muscle excitation (**Fig. 2A,B**). Together, selective silencing of MN-Ib and -Is inputs reveals chronic and acute PHP are expressed with opposing input specificity: Chronic PHP induces enhanced release at weak (low P_r_) MN-Ib synapses, while acute PHP drives enhanced release only at strong (high P_r_) MN-Is terminals.

**Figure 2:**
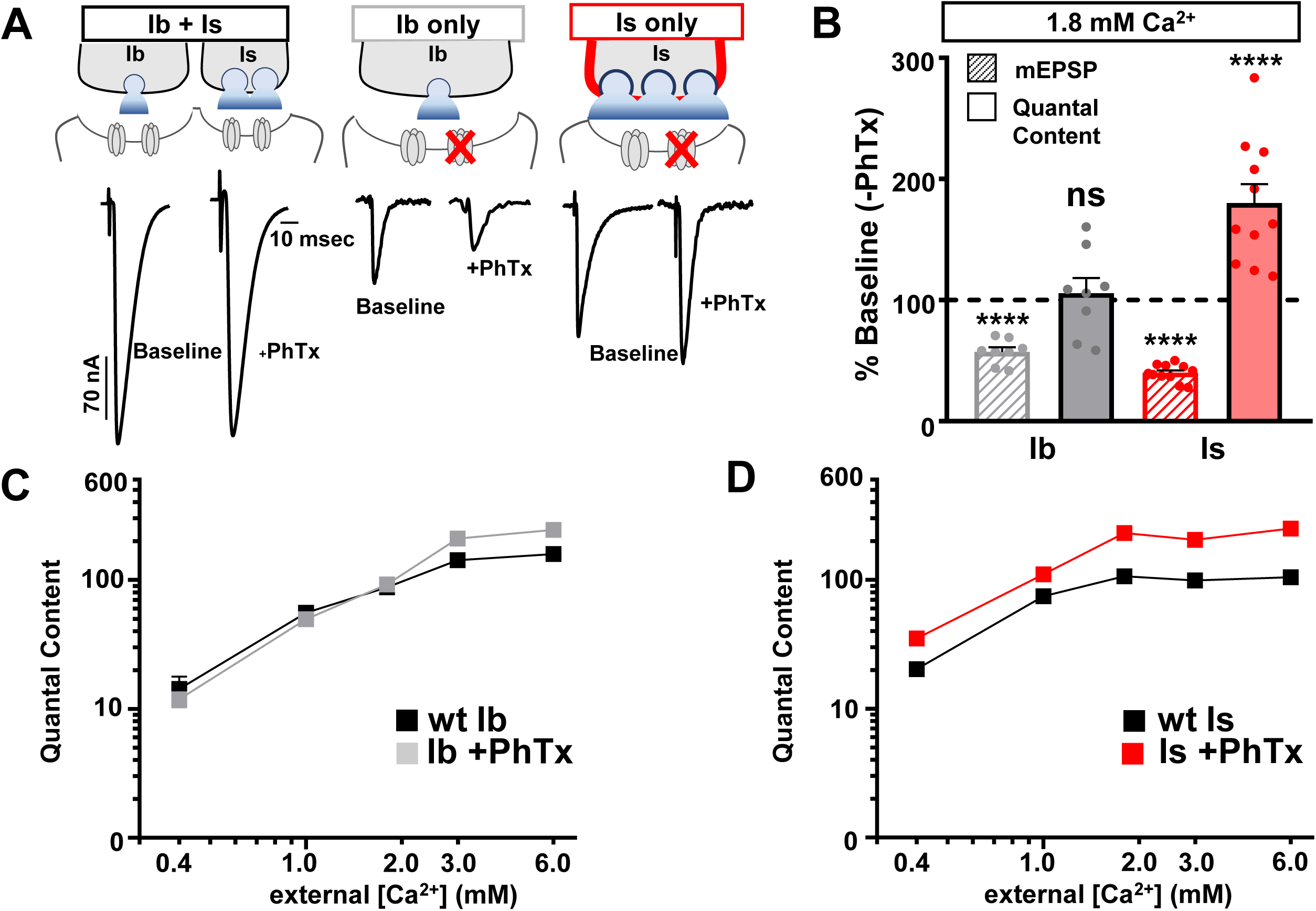
Acute PHP is selectively expressed at phasic MN-Is synapses. **(A)** Schematics and representative EPSC traces for baseline (wild type; *w^1118^*) and acute PHP (*w^1118^+*PhTx) NMJs at isolated MN-Ib and -Is inputs following selective expression of BoNT-C at 1.8 mM extracellular [Ca^2+^]. **(B)** Quantification of mEPSP amplitude and quantal content at MN-Ib and - Is after PhTx application normalized to baseline values. Note that while quantal content is enhanced at MN-Is, characteristic of robust PHP expression, no change is observed at MN-Ib. **(C,D)** Plot of quantal content as a function of external [Ca^2+^] for MN-Ib and -Is at baseline (wild type) and after acute PHP signaling (+PhTx) at MN-Ib (C) and MN-Is (D). Note that the enhanced quantal content characteristic of PHP saturates at 1.8 mM Ca^2+^ and above at MN-Is, while quantal content does not increase at MN-Ib at physiological Ca^2+^ and below. Error bars indicate ± SEM. Additional statistical details are shown in Table S1.

### Distinct remodeling of active zone nanostructure following input-specific PHP signaling

Next, we examined active zone structures at MN-Ib and -Is in wild type and after chronic vs acute PHP. Previous studies have shown that active zones remodel at MN-Ib after PHP signaling, with an apparent increase in fluorescence intensity of active zone components observed(Weyhersmuller et al., 2011; Goel et al., 2017; Li et al., 2018a; Böhme et al., 2019; Goel et al., 2019b; Gratz et al., 2019; Mrestani et al., 2021). In these experiments and for the rest of our study, we focused on changes at MN-Ib in *GluRIIA* mutants and MN-Is after PhTx compared to wild type controls at each input, as these are the relevant conditions for input-specific PHP expression. First, we imaged three components of the active zone using confocal microscopy: The core active zone scaffolds Bruchpilot (BRP) and Rim Binding Protein (RBP), as well as the Ca_v_2 voltage-gated Ca^2+^ channel subunit Cacophony (CAC). Similar to previous studies, we observed the intensity of all three to be enhanced after chronic PHP at MN-Ib terminals (**Fig. 3A,B**). Similarly, we observed that the intensity of these components were increased at MN-Is following acute PHP (**Fig. 3C,D**), consistent with a recent study(Medeiros et al., 2023). An increase in fluorescence intensity could indicate an increase in the abundance of the protein, as has been suggested to occur at MN-Ib after chronic PHP induction(Böhme et al., 2019; Goel et al., 2019b). However, an apparent increase in mean fluorescence intensity using confocal microscopy might instead reflect an increase in the density of the structure (“compaction”), which has been noted after PHP using super resolution approaches(Mrestani et al., 2021; Ghelani et al., 2023; Mrestani et al., 2023). These possibilities are not mutually exclusive. Hence, we employed super resolution <u>St</u>imulated <u>E</u>mission <u>D</u>epletion (STED) microscopy to understand how active zone components are remodeled after PHP signaling at tonic vs phasic terminals.

**Figure 3:**
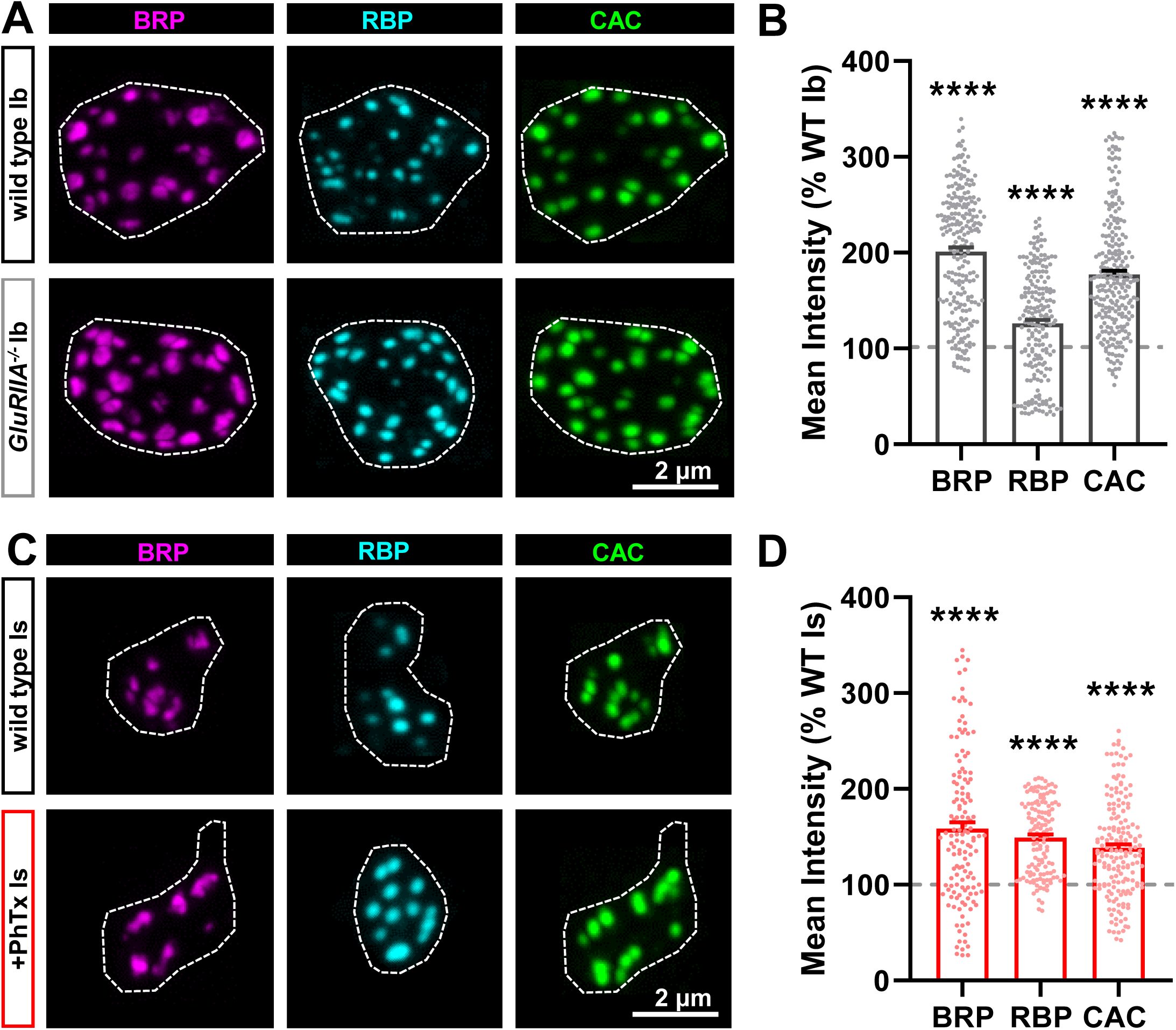
Active zone components remodel at both MN-Ib and -Is following PHP signaling. **(A)** Representative images of NMJs stained with anti-BRP, anti-RBP, and anti-CAC (GFP) at MN-Ib terminal boutons at muscle 6 of wild type (*cac^sfGFP^*) and chronic PHP (*cac^sfGFP^*;*GluRIIA^pv3^*). Dashed lines indicate the bouton boundary defined by HRP. **(B)** Quantification of mean fluorescence intensity in chronic PHP normalized to control (*cac^sfGFP^*). **(C,D)** Representative images and quantification as in (A,B) at MN-Is baseline (*cac^sfGFP^*) and after acute PHP signaling (*cac^sfGFP^*+PhTx). Error bars indicate ± SEM. Additional statistical details are shown in Table S1.

STED microcopy of BRP and CAC at MN-Ib revealed an increase in the area of both components after chronic PHP (**Fig. 4A,C**), with a corresponding enhancement in the number of BRP “nano-modules” (**Fig. 4B**), nodes of local maximal intensity(Böhme et al., 2019; Goel et al., 2019b; Muttathukunnel et al., 2022). We also observed an increase in the number of “merged” BRP rings at MN-Ib after chronic PHP signaling (**Fig. S1**), which has been previously reported(Hong et al., 2020). Merged BRP rings are associated with enlarged active zones with enhanced release probability(Graf et al., 2009; Goel et al., 2019b). However, no change in merged BRP rings was seen at MN-Is after acute PHP (**Fig. S1**). Area ratios of both CAC:BRP and RBP:BRP scaled in proportion (**Fig. 4D**), reflecting an apparent increase in the abundance of these active zone components spread across a larger area. In contrast, STED imaging of BRP and CAC at MN-Is after acute PHP revealed a compaction of the structures, with reduced CAC area (**Fig. 4G,I**) and no significant change in BRP nanomodules (**Fig. 4H**). Interestingly, while the Is areas of BRP and RBP did not significantly change after acute PHP, there was a selective reduction in the CAC area (**Fig. 4I,J**). These data suggest that active zone abundance does not change at MN-Is after acute PHP signaling; rather, the density of at least one component, CAC, is selectively increased, leading to a “compaction” of the Ca^2+^ channels(Ghelani et al., 2023). Thus, STED imaging reveals that chronic PHP expands active zone area and increases the abundance of material at MN-Ib release sites (schematized in **Fig. 4E,F**), while acute PHP compresses the density of at least CAC channels at MN-Is active zones without a change in protein abundance (schematized in **Fig. 4K,L**).

**Figure 4:**
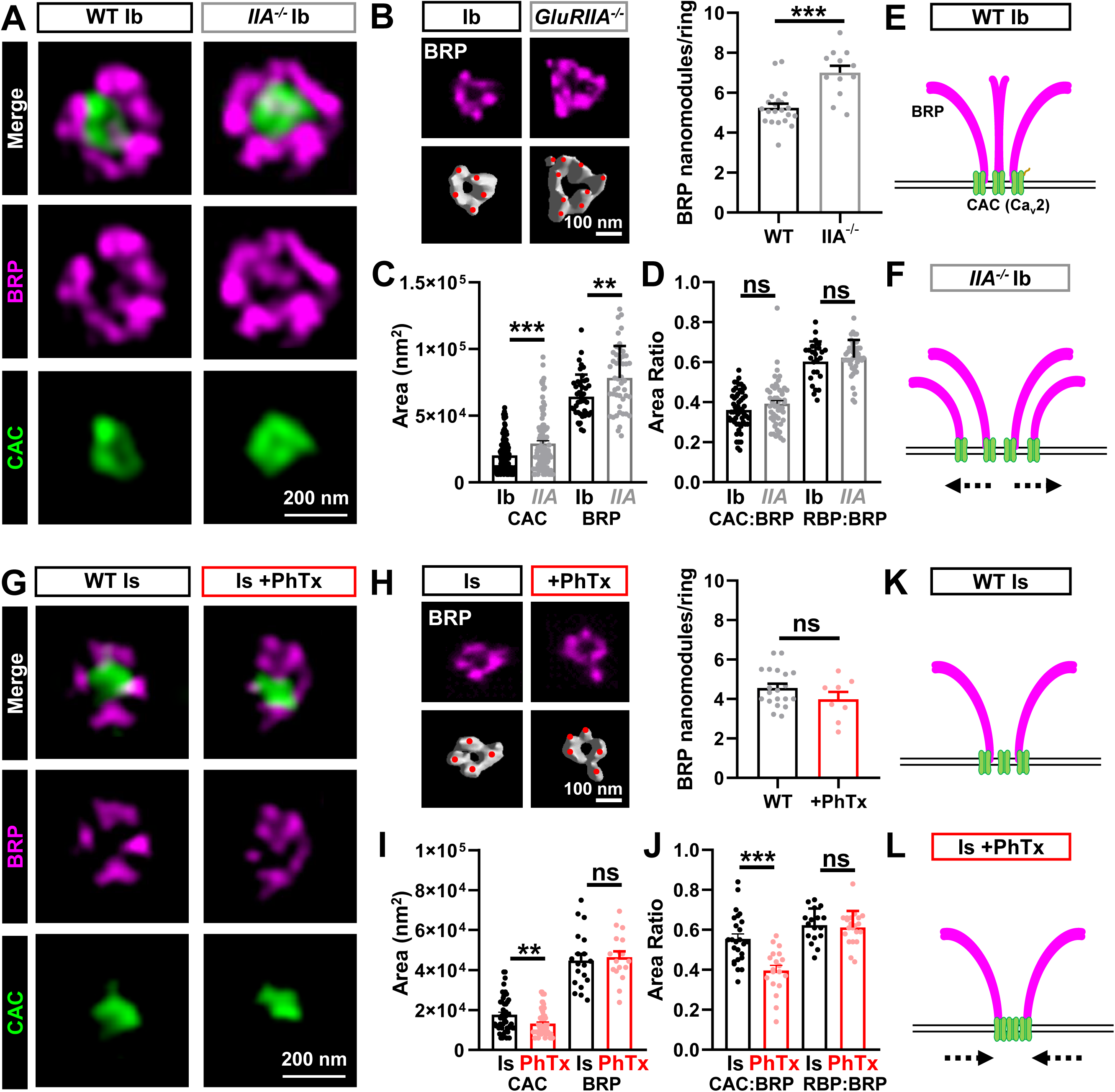
STED imaging reveals homeostatic expansion of active zones at MN-Ib and compaction at MN-Is. **(A)** Representative STED images of BRP and CAC at single MN-Ib boutons of wild type (*cac^sfGFP^*) and chronic PHP (*cac^sfGFP^;GluRIIA^pv3^*). **(B)** Representative images and quantification of BRP nanomodules in wild type and chronic PHP. **(C,D)** Quantification of CAC and BRP areas alone and area ratios in the indicated genotypes. Note that areas expand and scale together at MN-Ib after chronic PHP signaling. **(E,F)** Schematics showing homeostatic expansion of MN-Ib active zones following chronic PHP signaling. **(G)** Representative STED images of BRP and CAC at single MN-Is boutons at baseline and after PhTx application (acute PHP). **(H)** Representative images and quantification of BRP nanomodules in wild type and acute PHP. **(I,J)** Quantification of CAC and BRP areas alone and area ratios of the indicated conditions. Note that CAC puncta become more compact at MN-Is after acute PHP signaling. **(K,L)** Schematics indicating homeostatic compaction of MN-Is active zones following acute PHP signaling. Error bars indicate ± SEM. Additional statistical details are shown in Table S1.

### Input-specific PHP selectively targets presynaptic Ca^2+^ influx and vesicle pools

Enhanced abundance of CAC channels at MN-Ib after chronic PHP should increase presynaptic Ca^2+^ influx and promote neurotransmitter release, while acute PHP might alter CAC function to homeostatically tune release. To determine whether presynaptic Ca^2+^ levels change after PHP, we developed a ratiometric Ca^2+^ indicator, targeted to presynaptic boutons, using the highly sensitive genetically encoded Ca^2+^ indicator GCaMP8f(Zhang et al., 2023). Specifically, we fused the monomeric red-shifted fluorophore mScarlet(Bindels et al., 2017), which is not sensitive to Ca^2+^, to GCaMP8f(Li et al., 2021). To localize this indicator to synaptic boutons, we fused these proteins to the synaptic vesicle protein Synaptotagmin (Syt) to make **mScar8f** (Syt::mScarlet::GCaMP8f) (**Fig. 5A,D**). Using resonant area scans of single MN-Ib or -Is boutons, we confirmed the >two-fold larger baseline Ca^2+^ increase at MN-Is over Ib (**Fig. 5B,C,E,F**), previously reported using chemical dyes(Lu et al., 2016; He et al., 2023), which contributes to the strong P_r_ of MN-Is. We also found that chronic PHP increases the Ca^2+^ signal by ∼50% at Ib (**Fig. 5B,C**), consistent with previous reports using chemical dyes at MN-Ib(Muller and Davis, 2012). However, whether PHP changes Ca^2+^ levels at MN-Is has not been determined. mScar8f imaging of MN-Is revealed no significant change in the Ca^2+^ signal at MN-Is after PhTx application relative to baseline (**Fig. 5E,F**), nor in the rise or decay time constants (Table S1). Thus, chronic PHP is achieved at MN-Ib through a selective enhancement in presynaptic Ca^2+^ influx, while other mechanisms must be involved in acute PHP.

**Figure 5:**
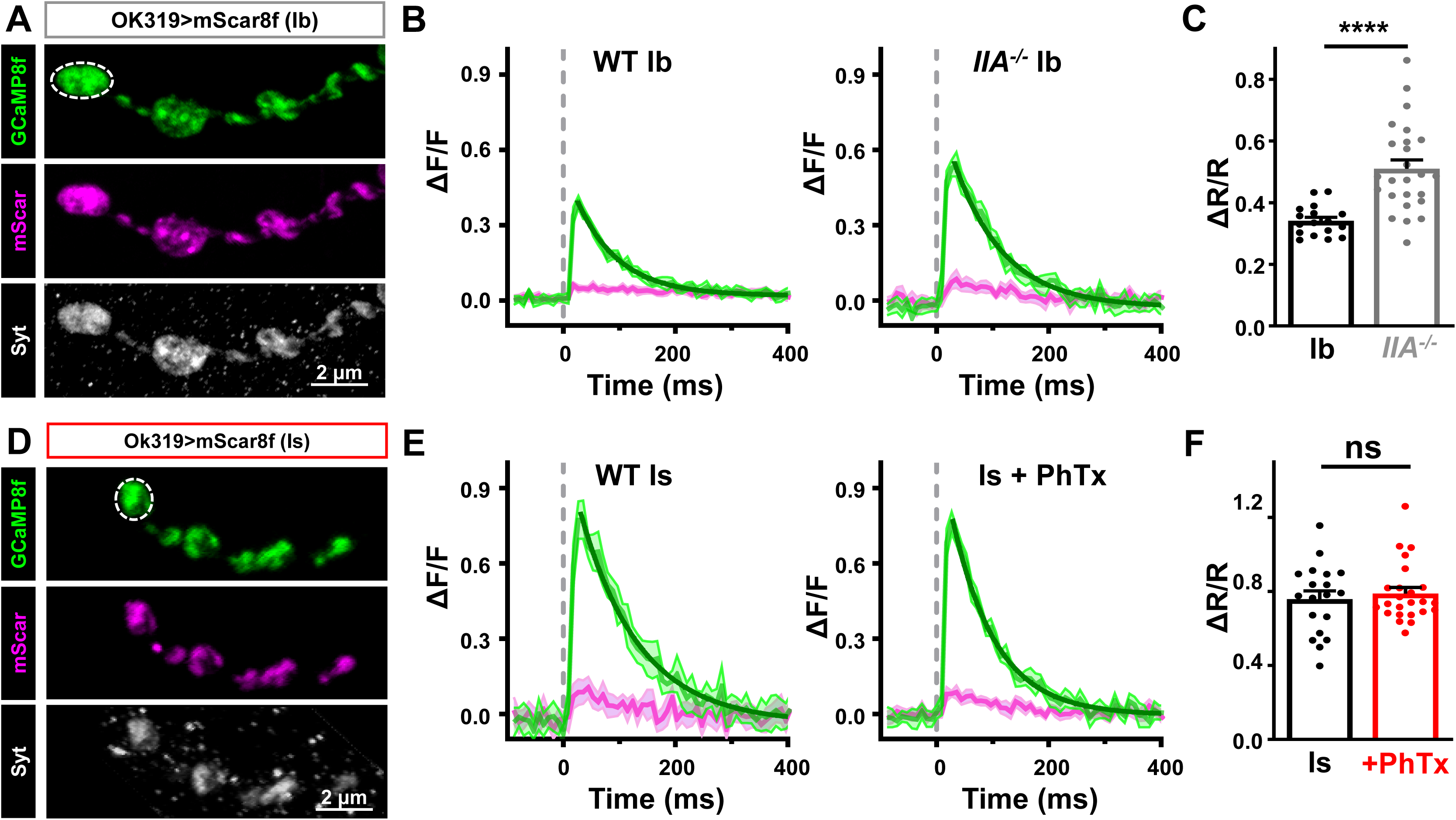
Acute PHP does not enhance presynaptic Ca^2+^ influx at phasic MN-Is terminals. **(A)** Representative immunostaining images of mScarlet and GCaMP8f at MN-Ib (OK319>UAS-Syt::mScarlet::GCamp8f) co-stained with anti-Syt. Note the bouton labeled with dashed lines represents the region of interest undergoing resonant area scanning. **(B)** Representative Ca^2+^ imaging traces from MN-Ib resonant area scans of mScar8f expressed in wild type and *GluRIIA* mutants. Averaged GCaMP8f (green) and mScarlet (magenta) signals from 10 stimuli; shadow indicates +/-SEM. Decays are fit with a one phase exponential (dark green). **(C)** Quantification of ΔR/R (GCaMP8f/mScarlet <u>R</u>atio) from MN-Ib boutons demonstrate that chronic PHP enhances Ca^2+^ signals, as expected. **(D-F)** Similar images, traces, and quantification as (A-C) but from mScar8f expression at MN-Is in wild type and acute PHP. Note that while baseline Ca^2+^ levels are higher at MN-Is boutons compared to MN-Ib, as expected, no change is observed after acute PHP signaling. Error bars indicate ± SEM. Additional statistical details are shown in Table S1.

The number of synaptic vesicles available for release, referred to as the *readily releasable vesicle pool* (RRP), has been shown to increase after PHP signaling(Weyhersmuller et al., 2011; Muller et al., 2012; Kiragasi et al., 2017; Li et al., 2018a). To determine the RRP size, synapses are stimulated at high frequency (60 Hz) at elevated P_r_ conditions (3 mM extracellular Ca^2+^), and the cumulative EPSC is plotted (**Fig. 6A,C**)(Li et al., 2018b). A linear line from stimulus 19-30 is fitted to time 0 (y-intercept) to estimate the cumulative EPSC, and each cumulative EPSC is normalized to its quantal size to estimate the size of the RRP(Li et al., 2018b) (see methods). Using this approach, we estimated the RRP at MN-Ib baseline and after chronic PHP. Surprisingly, we found no significant change in the RRP at MN-Ib following chronic PHP signaling (**Fig. 6A,B**). In contrast, the RRP was selectively increased at MN-Is after acute PHP, almost tripling in size (**Fig. 6C,D**). Thus, two distinct mechanisms are selectively targeted to achieve input-specific PHP: Chronic PHP enhances presynaptic Ca^2+^ influx at MN-Ib, while acute PHP increases the number of vesicles available for release at MN-Is terminals.

**Figure 6:**
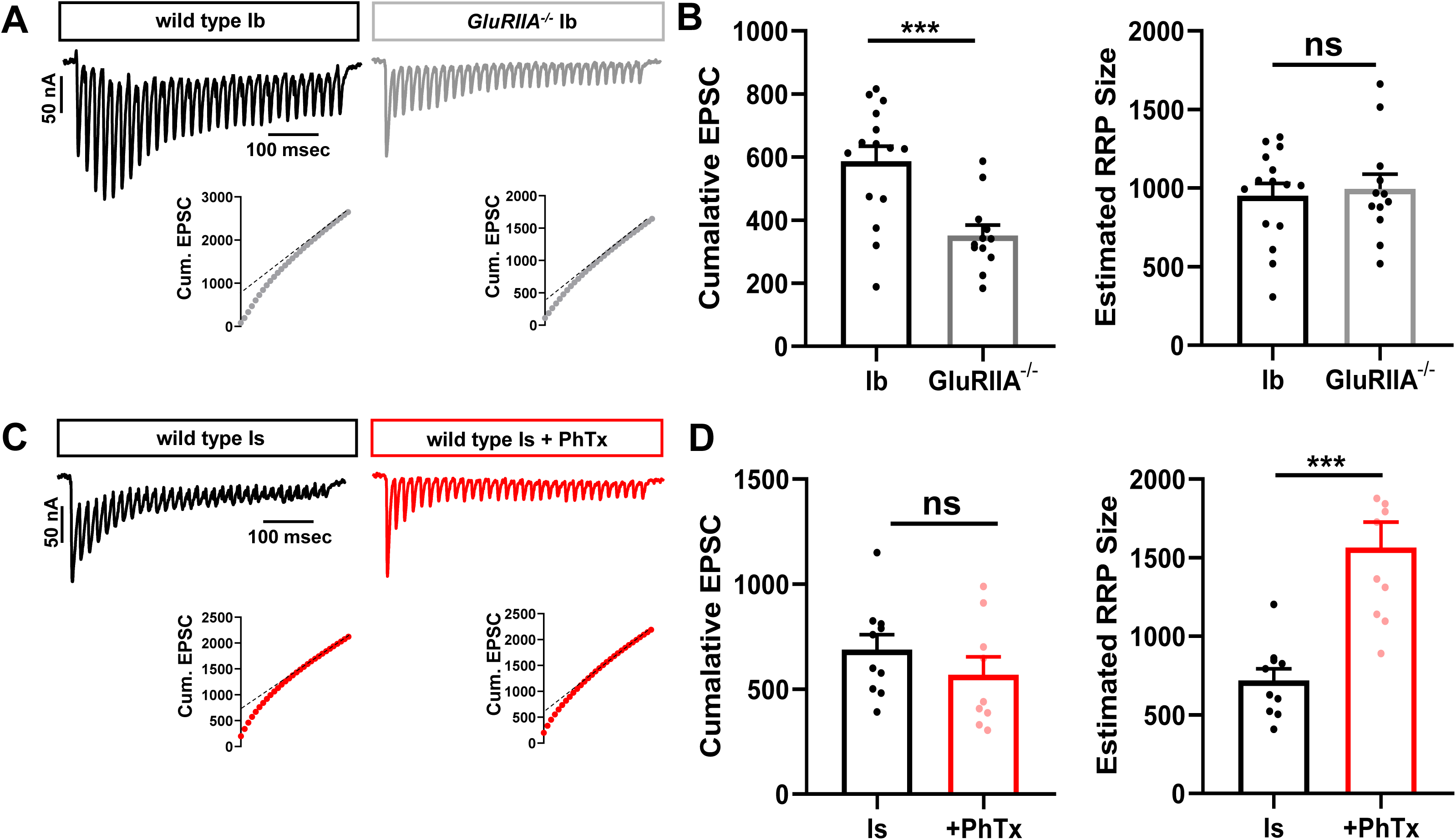
PHP selectively expands the RRP at phasic MN-Is synapses. **(A)** Representative traces of 30 EPSCs from wild type and *GluRIIA* mutant MN-Ib NMJs stimulated at 60 Hz in 3 mM extracellular [Ca^2+^]. Bottom: Averaged cumulative EPSC amplitudes. A line fit to the 18-30th stimuli was back extrapolated to time 0. **(B)** Quantification of cumulative EPSC and estimated RRP size at MN-Ib inputs of wild type and *GluRIIA* mutants. Note that chronic PHP does not change the RRP size at MN-Ib. **(C,D)** Representative traces and quantifications as shown in (A,B) from MN-Is of wild type and after PhTx application. Note that acute PHP enhances RRP size at MN-Is. Error bars indicate ± SEM. Additional statistical details are shown in Table S1.

### PHP targets functional release sites and vesicle coupling

In our final set of experiments, we examined two additional electrophysiological approaches, mean-variance analysis and Ca^2+^ coupling, to probe how PHP adaptations at tonic vs phasic synapses ultimately influence synaptic vesicle release properties. Mean-variance analysis is an approach that uses a mathematical equation, based on the probabilistic nature of vesicle fusion, to determine the number of release sites that function at a given synapse(Clements, 2003). In particular, the EPSC variance is plotted as a function of the average EPSC amplitude across a range of extracellular Ca^2+^ concentrations, with 0 variance observed at 0 and also at very high, saturating Ca^2+^ concentrations (see methods). From this analysis, one can estimate the number of functional release sites per entire NMJ, which previous studies have used to show that PHP enhances functional release site number at ambiguated NMJs (Weyhersmuller et al., 2011; Li et al., 2018b), likely through an Unc13-dependent mechanism(Böhme et al., 2016; Reddy-Alla et al., 2017; Ortega et al., 2018; Böhme et al., 2019; Jusyte et al., 2023). Using mean-variance analysis after isolating MN-Ib and -Is, we observed similar increases in the number of functional release sites at both MN-Ib and -Is after chronic and acute PHP (**Fig. 7A-C; G-I**), where the number of functional release sites was increased by 53% and 62%, respectively. To probe this enhancement in functional release sites in more detail, we examined Unc13. Unc13 is a fusogenic scaffold that positions synaptic vesicles for release at active zones(Jahn and Fasshauer, 2012; Reddy-Alla et al., 2017; Sakamoto et al., 2018; Dittman and Ryan, 2019), where activation of *Drosophila* Unc13A is thought to correlate with the number of functional release sites(Böhme et al., 2016; Reddy-Alla et al., 2017). Using confocal and STED microscopy, we observed an increase in the fluorescence intensity and area of the Unc13A signal at MN-Ib after chronic PHP (**Fig. 7D-F**), with similar changes observed at Is after acute PHP (**Fig. 7J-L**). These cell biological changes are thought to reflect increased activation of Unc13A, and are associated in an enhancement in the number of vesicle release sites at active zones(Böhme et al., 2016; Jusyte et al., 2023), ultimately converging on a need for Unc13A, which is necessary for both acute and chronic PHP expression(Böhme et al., 2019). Thus, both chronic and acute PHP targets Unc13A to enhance functional release site numbers and promote the homeostatic increase in presynaptic release.

**Figure 7:**
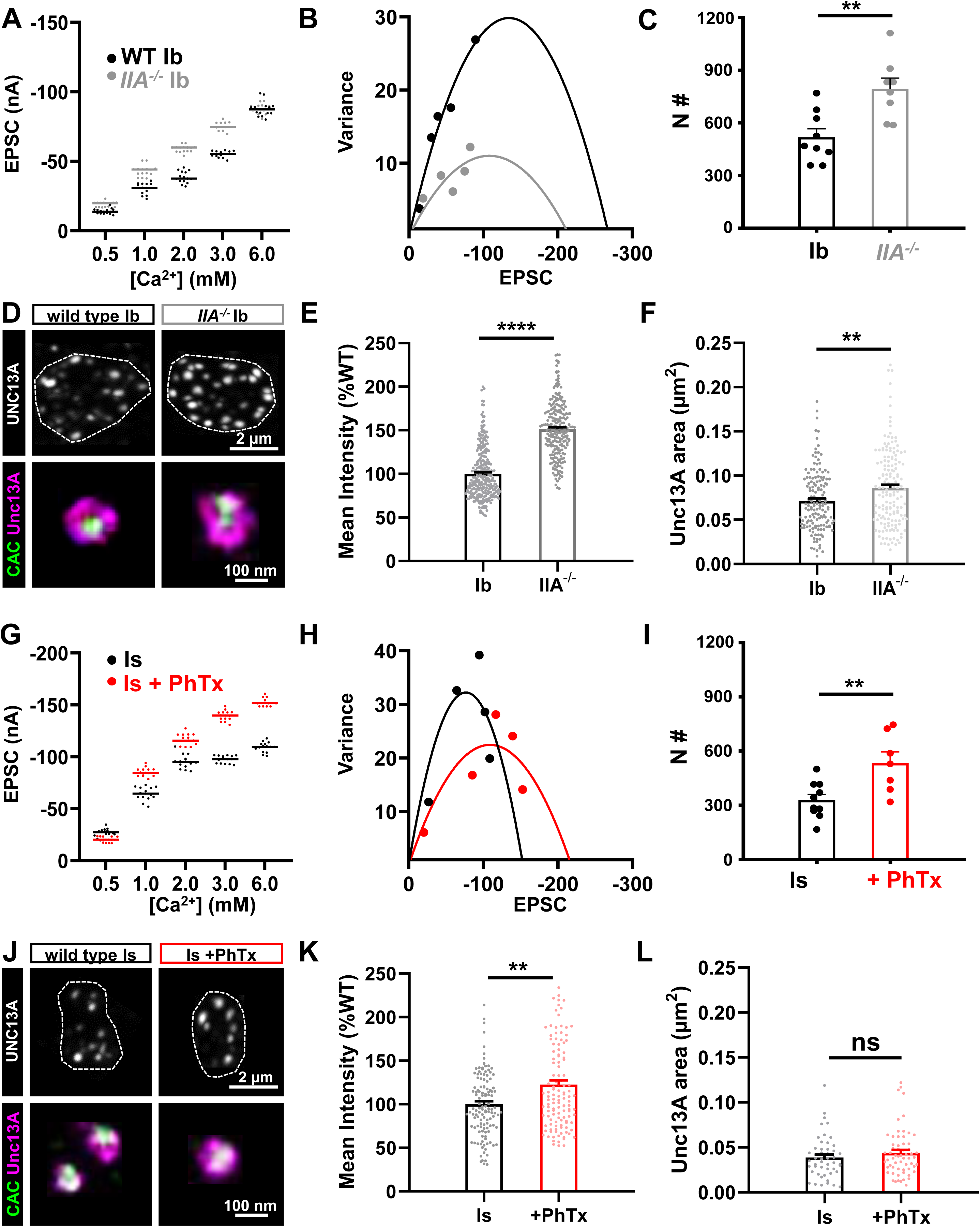
PHP increases functional release sites at both tonic and phasic inputs. **(A)** Representative scatter plot of EPSC amplitude distribution from MN-Ib NMJs in wild type and *GluRIIA* mutants in the indicated extracellular [Ca^2+^]. **(B)** Example mean-variance plot for the data shown in (A). Variance was plotted against the mean amplitude of 15 EPSCs from the five Ca^2+^ concentrations detailed in (A). **(C)** Estimated number of functional release sites based on mean-variance plots from multiple NMJ recordings, indicating enhanced release sites at MN-Ib after chronic PHP signaling. **(D)** Representative images of UNC13A and CAC immunostaining at MN-Ib boutons in the indicated genotypes using confocal and STED microscopy. Dashed lines indicate the neuronal membrane. **(E)** Quantification of UNC13A confocal mean fluorescence intensity in *GluRIIA* mutants normalized to wild type. **(F)** Quantification of UNC13A area before or after chronic PHP signaling. Note that both UNC13A area and intensity increase at MN-Ib following chronic PHP. **(G-L)** Same images and analyses as described in (A-F) at MN-Is boutons of wild type and following PhTx application. Note that acute PHP similarly enhances functional release sites at MN-Is inputs. Error bars indicate ± SEM. Additional statistical details are shown in Table S1.

Finally, we probed Ca^2+^ channel-vesicle coupling at tonic vs phasic synapses after PHP modulation. In this approach, the proportion of low P_r_ (“loosely coupled”) synaptic vesicles are estimated by competition for intracellular Ca^2+^ using the slow Ca^2+^ buffer EGTA(Meinrenken et al., 2002; Kaeser and Regehr, 2014). Under basal conditions, vesicles at the strong MN-Is are more tightly coupled compared to coupling at the weak MN-Ib(He et al., 2023). At MN-Ib NMJs, we observed a decrease in EGTA sensitivity after chronic PHP, suggesting a proportionate decrease in loosely coupled vesicles at these synapses, and a presumable shift in the proportion of tightly coupled vesicles (**Fig. 8A,B**). However, at MN-Is, we found an increase in EGTA sensitivity (**Fig. 8C,D**), suggesting an enhancement in the proportion of loosely coupled vesicles contributing to increased neurotransmitter release. Together, these data suggest an opposing, input-specific modulation of vesicle coupling during PHP: Proportionately fewer loosely coupled vesicles contribute to enhanced release after chronic PHP at MN-Ib, likely due to increased Ca^2+^ influx. In contrast, more loosely coupled vesicles are recruited to promote potentiation at MN-Is after acute PHP, likely due to compaction of active zone components and the addition of more release sites at the outer perimeter of active zones. We present a schematic summarizing the input-specific changes at tonic vs phasic release sites after PHP (**Fig. 8E**).

**Figure 8:**
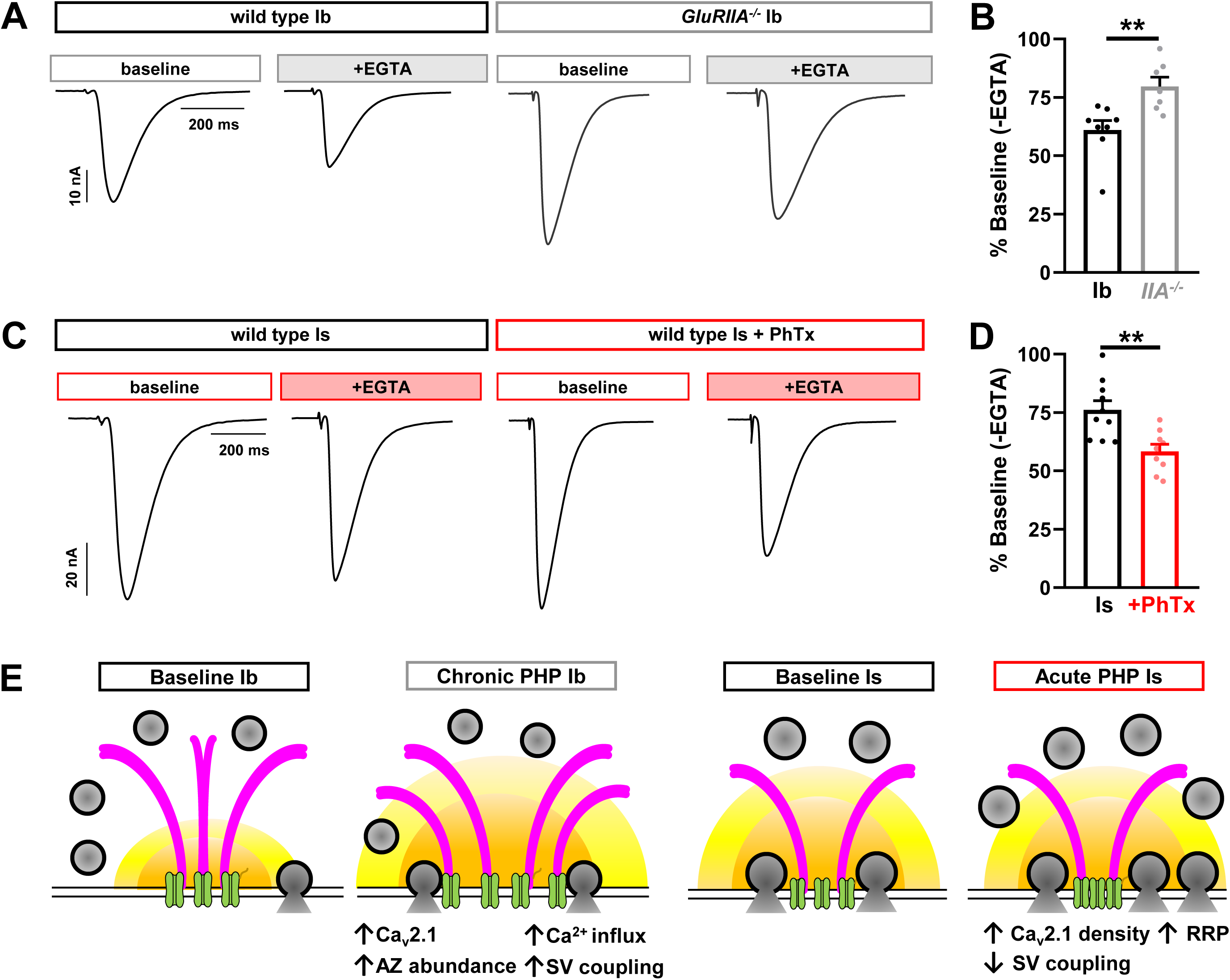
Chronic and acute PHP inversely modulate the loosely coupled synaptic vesicle pool. **(A)** Representative EPSC traces from isolated MN-Ib NMJs following incubation of EGTA-AM in wild type or *GluRIIA* mutants. **(B)** Quantification of EPSC amplitudes in wild type and *GluRIIA* mutants after EGTA-AM treatment normalized to baseline values. Note that there is an apparent decrease in the loosely coupled (EGTA sensitive) vesicle pool following chronic PHP. **(C,D)** Same traces and quantification as (A,B) but of wild type MN-Is NMJs at baseline and following PhTx application treated with EGTA-AM. Note that there is an apparent increase in the loosely coupled (EGTA-sensitive) vesicle pool. **(E)** Schematics summarizing input-specific mechanisms enabling chronic and acute PHP expression. Chronic PHP enhances Ca^2+^ influx at MN-Ib by enhancing active zone size and protein abundance, while acute PHP is achieved through compacting active zones and enhancing RRP vesicle pool size. Error bars indicate ± SEM. Additional statistical details are shown in Table S1.

## DISCUSSION

By electrophysiologically isolating transmission from tonic vs phasic motor inputs, we have illuminated the expression mechanisms that enable input-specific PHP expression. At physiological Ca^2+^ conditions and below, chronic and acute PHP selectively target distinct motor neuron inputs: Chronic PHP only enhances presynaptic release at tonic MN-Ib inputs, while acute PHP selectively potentiates phasic MN-Is neurons. Importantly, this selective modulation also targets distinct processes: Chronic PHP expands active zone nanostructures at tonic terminals, leading to enhanced presynaptic Ca^2+^ influx, more functional release sites, and more tightly coupled vesicles. In contrast, acute PHP contracts active zones and expands the readily releasable synaptic vesicle pool while engaging additional loosely coupled vesicles. Together, these findings resolve long-standing questions about whether and how presynaptic homeostatic plasticity selectively adjusts release at distinct synaptic inputs, while raising new conundrums about the trans-synaptic dialogue orchestrating PHP induction.

### Tonic motor neurons and chronic PHP

Chronic PHP homeostatically modulates presynaptic neurotransmitter release selectively at tonic motor neurons by sculpting the basal characteristics of these synapses. Tonic MN-Ib inputs function with low release probability, facilitating with high frequency stimulation, and engage a relatively large pool of loosely coupled synaptic vesicles(Lu et al., 2016; Newman et al., 2017; Aponte-Santiago et al., 2020; He et al., 2023; Medeiros et al., 2023). Indeed, active zones at MN-Ib terminals appear to be quite plastic, not only expanding in size and nanomodularity following chronic PHP signaling, but incorporating enhanced abundance of Ca_v_2 Ca^2+^ channels and scaffolds including BRP, RBP, and Unc13A(Goel et al., 2017; Böhme et al., 2019; Goel et al., 2019b; Gratz et al., 2019). Protein transport is essential for the expression of chronic PHP, as indicated by the requirement of the molecular motors Aplip-1, Srpk79D, and Arl8 in the transport of this cargo during PHP at MN-Ib presynaptic compartments(Böhme et al., 2019; Goel et al., 2019b). Notably, tonic motor inputs innervate individual muscle fibers to drive contraction through summation of repetitive stimulation(Newman et al., 2017), and it seems likely that this property must be maintained *in vivo* following chronic PHP signaling to ensure proper muscle activity. Hence, while we only assessed synaptic strength through single action potential stimulation, the attributes of PHP at tonic inputs likely serve to stabilize muscle excitability through maintaining basal patterns of activity.

It is interesting to note that while chronic PHP is selectively expressed at tonic inputs at physiological Ca^2+^ and below (0.4-1.8 mM), both tonic and phasic neurons appear to release enhanced neurotransmitter following loss of *GluRIIA* at high Ca^2+^ conditions (3-6 mM; **Fig. 1C,D**), as previously reported(Genç and Davis, 2019). The apparent enhancement of release at phasic MN-Is in *GluRIIA* mutants, and the physiological relevance of this change, is difficult to interpret. First, no functional change is observed at MN-Is in physiological Ca^2+^ and below (1.8 mM) despite loss of *GluRIIA*. Second, at baseline states, MN-Is neurotransmitter release saturates above 1.8 mM Ca^2+^(He et al., 2023). Nonetheless, the apparent enhancement of release at elevated Ca^2+^ conditions at both tonic and phasic inputs raises questions about whether loss of *GluRIIA* leads to a selective induction of PHP at tonic MN-Ib, or, alternatively, whether chronic PHP is induced at both inputs, but the functional enhancement at phasic neurons is only observed at highly elevated, non-physiologic Ca^2+^ conditions.

How chronic PHP is induced remains enigmatic, although it does not depend on reduced postsynaptic Ca^2+^ influx(Goel et al., 2017; Perry et al., 2022). Rather, CaMKII activity and the GluRIIA C-tail appear to be intimately involved in chronic PHP induction(Perry et al., 2022), where active CaMKII is selectively enriched at postsynaptic compartments of tonic MN-Ib NMJs(Newman et al., 2017; Li et al., 2018b; Perry et al., 2022). It is also important to highlight that chronic PHP can be induced with apparent synapse specificity: Loss of *GluRIIA* at a single muscle can selectively induce chronic PHP at the presynaptic release sites innervating that muscle without altering release from neighboring active zones of the same neuron innervating an adjacent muscle with normal glutamate receptor levels(Li et al., 2018b). Hence, chronic PHP can be induced and expressed with both target- and input-specificity.

### Phasic motor neurons and acute PHP

While imaging studies examining PHP-dependent changes at large tonic MN-Ib boutons have been extensively reported, far less was known about how the small phasic MN-Is terminals adjust to PHP signaling. We found that many of the homeostatic adaptations at MN-Ib boutons – namely enhanced active zone scaffolds, Ca^2+^ channel abundance, and Ca^2+^ influx – do not happen at phasic MN-Is terminals. Instead, Ca^2+^ channel nanostructure becomes more dense by STED microscopy, a structural remodeling also reported using other super resolution imaging modalities(Mrestani et al., 2021; Dannhäuser et al., 2022; Ghelani et al., 2023; Mrestani et al., 2023). While Ca^2+^ influx does not change at phasic terminals, the compaction of active zones likely drives the key homeostatic adaptation at phasic release sites - more synaptic vesicles available for release. There are a higher proportion of loosely-coupled vesicles at phasic terminals after acute PHP signaling, which likely responds to the same Ca^2+^ levels to enable enhanced glutamate emission. Thus, compaction of Ca^2+^ channels may enable more loosely-coupled vesicles to position for release along with Unc13A activation, motifs that tune release probability at a variety of synapse types(Rebola et al., 2019; Jusyte et al., 2023).

Previous studies have found that acute PHP remodels tonic MN-Ib active zones similarly to chronic PHP, so it is surprising how selective acute PHP is in functionally targeting only the phasic MN-Is input for potentiation after PhTx application. While acute PHP homeostatically potentiates presynaptic release from phasic MN-Is inputs across all Ca^2+^ conditions assayed, no change in quantal content was observed at MN-Ib across physiological Ca^2+^ conditions and below (**Fig. 2C**). Acute PHP has been reported in many previous studies to remodel active zone components at tonic MN-Ib, including BRP and CAC, similarly to what is seen in chronic PHP(Weyhersmuller et al., 2011; Goel et al., 2017; Böhme et al., 2019; Goel et al., 2019b; Gratz et al., 2019; Mrestani et al., 2021; Medeiros et al., 2023). Furthermore, PhTx application induces reorganization of postsynaptic glutamate receptors at MN-Ib NMJs in nano-alignment with presynaptic active zone structures(Muttathukunnel et al., 2022). Finally, acute PHP was reported to enhance Ca^2+^ influx at tonic Ib terminals(Muller and Davis, 2012). However, while these structural and imaging changes might seem to be shared similarly between acute and chronic PHP at MN-Ib, they appear to be functionally silent for acute PHP, as they lead to no significant change in presynaptic release. Consistent with this finding, a recent study found that when CAC remodeling is blocked, acute PHP is still robustly expressed(Ghelani et al., 2023). Thus, changes in active zone structure and Ca^2+^ levels during PHP do not necessarily have functional impacts, at least for single action potential stimulation at MN-Ib. The reasons for this are unclear, although ultrastructural organization of synaptic vesicles, Ca^2+^ channels, and Unc13 can collaborate to reduce release probability despite enhanced Ca^2+^ influx at Granule cell-Purkinje synapses in rodents(Rebola et al., 2019).

A major question for future studies centers on why a seemingly similar diminishment in glutamate receptor function at both tonic and phasic NMJs leads to such selective differences in presynaptic strength at each input. More specifically, how does genetic loss vs pharmacological blockade of GluRIIA-containing receptors lead to the selective expression of PHP at tonic or phasic inputs? A number of genes have been identified that are necessary for chronic, but not acute, PHP, including *brp(Frank et al., 2009; Marie et al., 2010; Spring et al., 2016; Böhme et al., 2019; James et al., 2019)*. The reasons for such distinct genetic requirements for chronic vs acute PHP are unclear, but it is tempting to now hypothesize that they may have specialized roles for plasticity at tonic vs phasic motor inputs. Beyond genetic distinctions, another contributing factor might involve input-specific differences in glutamate receptors and associated factors. While both GluRIIA- and GluRIIB-containing receptors are present at NMJs of both tonic and phasic inputs, GluRIIA:GluRIIB ratios are higher at tonic Ib NMJs(DiAntonio et al., 1999; Han et al., 2022; Han et al., 2023). Although receptors at both NMJs are inhibited, diminished postsynaptic Ca^2+^ influx does not seem to be involved in either chronic or acute PHP induction(Goel et al., 2017; Perry et al., 2022). In addition, tonic MN-Ib NMJs exhibit elaborate subsynaptic reticulum (SSR) structures(Jia et al., 1993; Teodoro et al., 2013; Nguyen and Stewart, 2016), where CaMKII is particularly enriched(Koh et al., 1999; Perry et al., 2022).

Interestingly, disrupted SSR morphology at fly NMJs also inhibit chronic PHP expression(Koles et al., 2015). Beyond the SSR, acute PHP must utilize a distinct induction mechanism, since the GluRIIA C-tail remains present after pharmacological blockade. Notably, a recent study suggested that acute PHP is induced through non-ionic signaling(Nair et al., 2021). Much remains to be learned about how glutamate receptor loss vs pharmacological blockade enables distinct retrograde signaling and presynaptic reorganization to enable input-specific adaptive plasticity.

## MATERIALS AND METHODS

### Fly stocks

*Drosophila* stocks were raised at 25°C using standard molasses food. Unless otherwise specified, the *w^1118^* strain was used as the wild-type control as this is the genetic background in which all genotypes were bred. For input-specific silencing experiments, MN-Ib and MN-Is only larvae were generated by crossing UAS-BoNT-C with Is-GAL4 (GMR27E09-GAL4) or Ib-GAL4 (dHb9-GAL4) as described(Han et al., 2022). We should note that expression of BoNT-C with these relatively weak drivers does not induce toxicity in motor neurons through early third-instar larval stages. However, we have found that expression of BoNT-C with stronger motor neuron drivers and/or for longer periods can perturb or even kill neurons. Endogenously tagged Cac^sfGFP-N^(Gratz et al., 2019) was used to label CAC, and *GluRIIA^PV3^*(Han et al., 2023) mutant backgrounds were used to induce chronic PHP expression. All experiments were performed on *Drosophila* third-instar larvae of both sexes. See Table S2 (Key Resources Table) for a full list of all fly stocks, antibodies, software, and their sources used in this study.

### Electrophysiology

All dissections and two-electrode voltage clamp (TEVC) recordings were performed as described(Kikuma et al., 2019) in modified hemolymph-like saline (HL-3) containing (in mM): 70 NaCl, 5 KCl, 10 MgCl_2_, 10 NaHCO_3_, 115 Sucrose, 5 Trehelose, 5 HEPES, pH=7.2, and CaCl_2_ at the specified concentration. To acutely block postsynaptic receptors, semi-dissected larvae (dorsally cut open with internal guts and nervous system intact) were incubated with philanthotoxin-433 (PhTx, 20µM; Sigma) in HL-3 for 10 min(Kiragasi et al., 2017). Internal guts, brain and the ventral nerve cord were subsequently removed to acquire fully dissected preparations. For PhTx-induced acute PHP experiments, fully dissected samples were thoroughly washed with HL3 three times before recording.

Recordings were carried out on an Olympus BX61 WI microscope stage equipped with a 40x/0.8 NA water-dipping objective and acquired using an Axoclamp 900A amplifier (Molecular Devices). Data were acquired from cells with an initial resting potential between -60 and -75 mV, and input resistances >5 MΩ. All recordings were conducted on abdominal muscle 6, segment A3 of third-instar larvae. The mEPSPs for each sample were recorded for 60 secs and analyzed with MiniAnalysis (Synaptosoft) and Excel (Microsoft) software. The average mEPSP amplitude for each NMJ were obtained from ∼100 events in each recording. Excitatory postsynaptic currents (EPSCs) were recorded by delivering 20 electrical stimuli at 0.5 Hz with 0.5 msec duration to motor neurons using an ISO-Flex stimulus isolator (A.M.P.I.) with stimulus intensities set to avoid multiple EPSCs.

The size of the readily releasable pool (RRP) was estimated as described(Goel et al., 2019a). Specifically, EPSCs were evoked with a 60 Hz, 30 stimulus train while recording in HL-3 supplemented with 3 mM Ca^2+^. The cumulative EPSC data was used to fit a line to the linear phase (stimuli #18–30) and back-extrapolated to time 0. The RRP was estimated by dividing the extrapolated EPSC value at time 0 by the average mEPSP amplitude. For the mean-variance plot, data was obtained from TEVC recordings using an initial 0.5 mM Ca^2+^ concentration, which was later increased to 1.0, 1.8, 3.0, and 6.0 mM through saline exchange via a peristaltic pump (Langer Instruments, BT100-2J) as described(He et al., 2023). EPSC amplitudes were monitored during the exchange, and 15 EPSC recordings (0.3 Hz stimulation rate) were performed in each condition. The variance (squared standard deviation) and mean (averaged evoked amplitude) were calculated from the 15 EPSCs at each individual Ca^2+^ concentration. GraphPad Prism was used to plot the variance against the mean for each concentration, with a theoretical data point at 0 variance and mean for Ca^2+^-free saline. Data from these six conditions were fit with a standard parabola (variance=Q*Ī -Ī2/N), where Q is the quantal size, Ī is the mean evoked amplitude (x-axis), and N is the functional number of release sites. N, as a parameter of the standard parabola, was directly calculated for each cell by the best parabolic fit.

To measure EGTA sensitivity, larval fillets were incubated in 0 Ca^2+^ modified HL-3 supplemented with 50 μM EGTA-AM (Sigma-Aldrich) for 10 min, then washed with HL-3 three times before recording in standard saline. EGTA-AM was applied following 10 mins PhTx incubation where applicable.

### Immunocytochemistry

Third-instar larvae were dissected in ice cold 0 Ca^2+^ HL-3 and immunostained as described(Chen et al., 2017; Kikuma et al., 2017; Perry et al., 2017). Briefly, larvae were either fixed in 100% ice-cold methanol for 5 min or 4% paraformaldehyde (PFA) for 10 mins followed by washing with PBS containing 0.1% Triton X-100 (PBST) for 10 mins, three times. Samples were blocked with 5% Normal Donkey Serum and incubated with primary antibodies overnight at 4°C. Preparations were washed for 10 mins thrice in PBST, incubated with secondary antibodies for 2 hours at room temperature, washed thrice again in PBST, and equilibrated in 70% glycerol. Prior to imaging, samples were mounted either in VectaShield (Vector Laboratories, for confocal) or ProLong Glass Antifade Mountant (ThermoFisher Scientific, for STED). For confocal experiments, native CAC-GFP was imaged. Other antigens were detected using the following primary antibodies: Mouse anti-BRP (nc82; 1:200); chicken anti-GFP (1:400); guinea pig anti-RBP (1:2000); guinea pig anti-Unc13A (1:500); Alexa Fluor 647-conjugated goat anti-Horseradish Peroxidase (HRP; 1:400). Secondary antibodies: STAR RED-conjugated secondary antibodies (1:200) were used for imaging in the infrared STED channel, while the others (Cy3-, AF488-, AF594- and AF647-conjugated) were used at 1:400. See Table S2 for a full list of all antibodies used and their sources.

We found that PhTx-induced remodeling effects were variable using standard PhTx treatments. We therefore developed the following protocol: For imaging experiments, PhTx was used at twice the concentration typically used for electrophysiology (40µM). PhTx was applied to semi-intact preps (dorsal incision only) held in place by magnetic pins so as not to perturb the body wall more than necessary. The tissue was allowed to incubate in PhTx at room temp for 15 mins before the dissection was completed. Fillets were then transferred to a standard dissection plate, stretched, pinned and fixed. Whenever possible, preparations were stained with BRP, and consistent remodeling was confirmed by an increase in BRP intensity at MN-Ib terminals.

### Confocal imaging and analysis

Confocal images were acquired with a Nikon A1R Confocal microscope equipped with NIS Elements software and a 100x APO 1.40NA oil immersion objective using separate channels with four laser lines (405 nm, 488 nm, 561 nm, and 647 nm) as described(Kiragasi et al., 2020). For fluorescence intensity quantifications of BRP, RBP, and CAC, z-stacks were obtained on the same day using identical gain and laser power settings with z-axis spacing of 0.150 µm and x/y pixel size of 40nm for all samples within an individual experiment. Raw confocal images were deconvolved with SVI Huygens Essential 22.10 using built-in Express settings. The default settings in SVI Huygen’s object analyzer were used to identify individual puncta within the 3D rendering and determine mean intensity of each punctum. All measurements based on confocal images were taken from M6/7 terminal boutons (1 bouton/Ib; 1-3 boutons/Is) acquired from at least 10 NMJs from four different animals.

### STED imaging and analysis

Stimulated Emission Depletion (STED) super resolution microscopy was performed as described(He et al., 2023). Briefly, STED imaging was performed using an Abberior STEDYCON system mounted on a Nikon Eclipse FN1 upright microscope equipped with four excitation lasers (640, 561, 488, and 405 nm), a pulsed STED laser at 775 nm, and three avalanche photodiode detectors that operate in a single photon counting mode. Multichannel 2D STED images were acquired using a 100x Nikon Plan APO 1.45 NA oil immersion objective with 15 nm fixed pixel size and 10 µsec dwell time using 15x line accumulation in photon counting mode and field of view of 1-2 boutons. Two secondary dyes were used, Abberior STAR Red and Alexa Fluor 594, and were depleted at 775 nm. For the STED channel, time gating was set at 1 nsec with a width of 6 nsec for all channels. Fluorescence photons were counted sequentially by pixel with the respective avalanche photodiode detector (STAR RED: 675±25 nm, Alexa Fluor 594: 600±25 nm). Raw STED images were deconvolved with the SVI Huygens software using the default settings and theoretical point spread functions for STED microscope. Covered areas of each protein were based on the raw STED images in red and infrared channels and quantified with the general analysis toolkit of NIS Elements software (Version 4.2). Active zones with optimal planar orientation were manually selected and the area and equivalent diameter of each protein was determined by applying an intensity threshold to mask layers in 600 nm and 675 nm channels, and a fixed intensity threshold value was applied to the same channels across samples. The nanomodules of each BRP puncta were quantified using local maxima detection in ImageJ with the same settings for all images. All measurements were based on 2D STED images that were taken from NMJs of M6 in segments A3/4 acquired from at least four different animals.

### Ca^2+^ imaging and analysis

Live presynaptic Ca^2+^ imaging was conducted in third-instar larvae NMJs expressing mScar8f (OK319>UAS-Syt::mScarlet::GCamp8f) as detailed(Chen et al., 2024). In brief, dissected third-instar larvae were immersed in HL3 containing 1.8mM Ca^2+^ for live imaging. 15 sec timelapse videos of terminal boutons from MN-Ib or -Is were acquired at 113 frames per second by a Nikon Eclipse Ni-E upright microscope equipped with a 60x 1.0NA water immersion objective and resonant scanner. The GCaMP8f signal was captured through the FITC channel (488nm excitation), while the mScarlet signal was recorded via the TRITC channel (561nm excitation). Electrical stimulation of the motor neuron was performed at 1 Hz with a 1 msec duration for the entire imaging session. The change in GCaMP8f and mScarlet mean intensities across frames was quantified using NIS Elements software. To correct for potential artifacts caused by muscle contraction, the GCaMP8f/mScarlet intensity ratio (R) of each frame was calculated to determine the Ca^2+^ influx-induced response. The response amplitude of each terminal was determined as the average of at least 10 stably recorded events. All measurements were based on the most terminal motor neuron boutons at M6 of segments A3 and A4.

### Statistical analysis

Data were analyzed using GraphPad Prism (version 8.0), MiniAnalysis (Synaptosoft), SVI Hugyens Essential (Version 22.10), or Microsoft Excel software (version 16.22). Sample values were tested for normality using the D’Agostino & Pearson omnibus normality test which determined that the assumption of normality of the sample distribution was not violated. Data were then compared using either a one-way ANOVA and tested for significance using a Tukey’s multiple comparison test or using an unpaired/paired 2-tailed Student’s t-test with Welch’s correction. In all figures, error bars indicate ±SEM, with the following statistical significance: p<0.05 (*), p<0.01 (**), p<0.001 (***), p<0.0001 (****); ns=not significant. Additional statistical details for all experiments are summarized in Table 1.

## Supporting information

Key Resources Table

## AUTHOR CONTRIBUTIONS

C.C., K.H., and D.D. designed the research; C.C, K.H., S.P., E.T., Y.H., and X.L. performed experiments and analyzed the data. The manuscript was written by C.C., K.H., and D.D. with feedback from the other authors.

## ACKNOWLEDGEMENTS

We thank Igor Delvendahl (University of Zurich, Switzerland) for important discussions on vesicle pool analyses. We acknowledge the Developmental Studies Hybridoma Bank (Iowa, USA) for antibodies used in this study and the Bloomington Drosophila Stock Center for fly stocks (NIH P40OD018537). This work was supported by grants from the National Institutes of Health to D.D (NS091546 and NS126654).

---

**Supplemental Figure 1:**
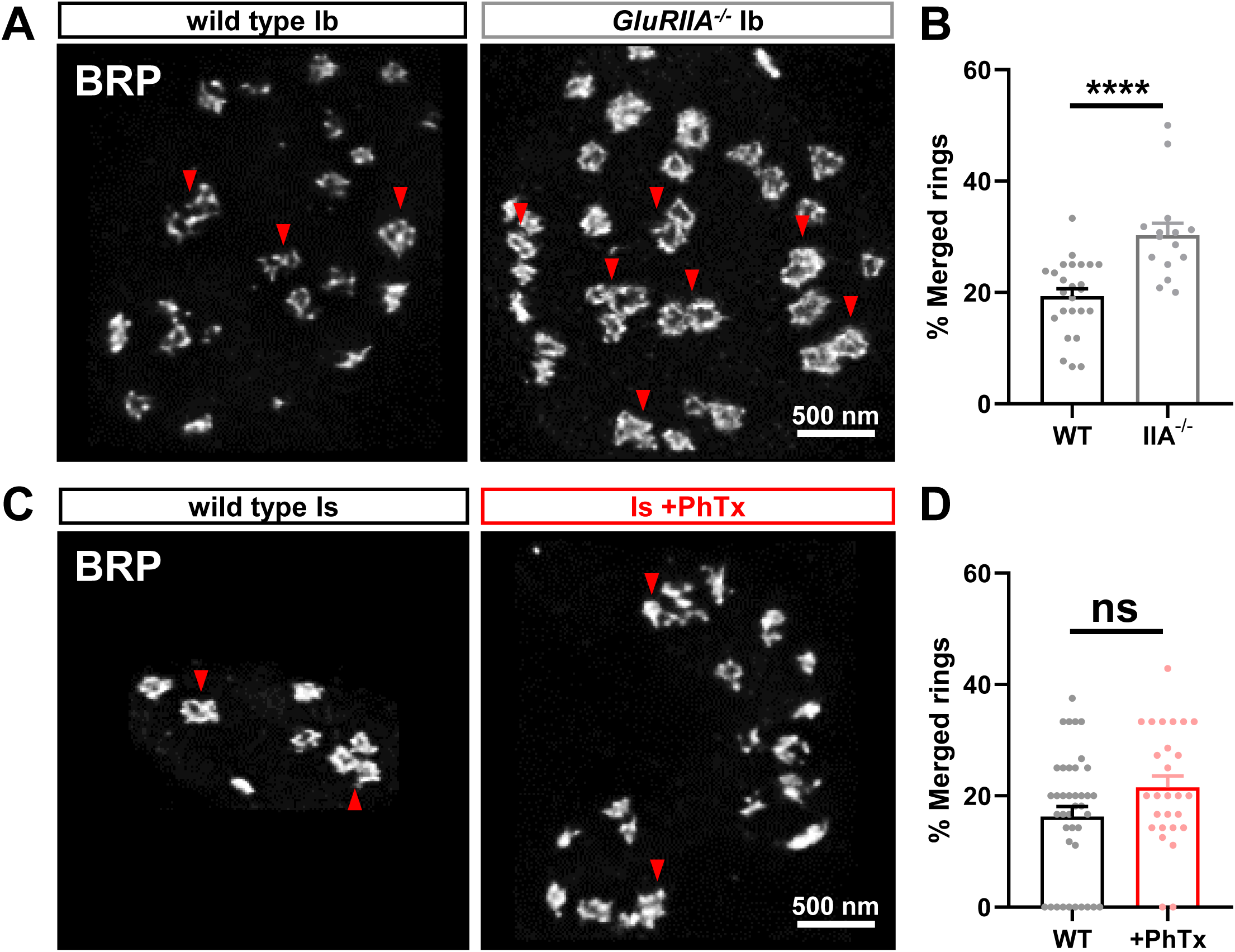
PHP signaling does not induce merging of BRP rings at phasic MN-Is synapses. **(A)** Representative STED images of BRP at MN-Ib terminal boutons in the indicated genotypes. **(B)** Quantification of merged BRP rings at MN-Ib at baseline of chronic PHP. MN-Ib has increased merged BRP rings following chronic PHP signaling. **(C,D)** Similar images and quantification as (A,B) in MN-Is terminal boutons before or after PhTx application. No significant change in merged BRPs rights is observed.

**Supplemental Table 1: Absolute values for normalized data and additional statistical details.** The figure and panel, genotype, extracellular Ca^2+^ concentration and conditions are noted. Average values (with standard error of the mean noted in parentheses), data samples (n), and statistical significance tests are shown for all data.

**Supplemental Table 2: Key resources table.**

