## Supplemental Table 1 for "Distinct input-specific mechanisms enable presynaptic homeostatic plasticity"

| Figure | Label | $[Ca^{2+}]$ (mM) | Genotype | mEPSP amp (mV) | EPSC amp (-nA) | QC | mEPSP freq.(Hz) | $R_{in}$ (MΩ) | $V_{rest}$ (mV) | n | One-way ANOVA, P Value: mEPSP, EPSC, QC. |
|---|---|---|---|---|---|---|---|---|---|---|---|
| 1A, B | wild type | 1.8 | *w*$^{1118}$ | 1.01 (±0.03) | 180.70 (±10.79) | 171.00 (±10.83) | 4.45 (±0.27) | 8.75 (±0.41) | -64.22 (±1.01) | 8 | - |
| 1A, B | GluRIIA | 1.8 | *GluRIIA*$^{pv3}$ | 0.42 (±0.03) | 155.6 (±13.28) | 380.96 (±40.72) | 1.80 (±0.36) | 7.75 (±0.56) | -75.00 (±2.29) | 7 | <0.0001 (****), 0.22 (ns), <0.0001 (****) |
| 1A, B | Ib only | 1.8 | *w;+;R27E09-GAL4/UAS-BoNT-C* | 0.59 (±0.06) | 52.35 (±4.15) | 87.60 (±5.90) | 1.98 (±0.22) | 9.78 (±0.62) | -72.86 (±1.81) | 11 | - |
| 1A, B | IIA, Ib only | 1.8 | *w;GluRIIA*$^{pv3}$*;R27E09-GAL4/UAS-BoNT-C* | 0.29 (±0.01) | 72.47 (±6.86) | 256.29 (±17.9) | 1.38(±0.10) | 10.11 (±0.53) | -67.51 (±2.03) | 8 | <0.0001 (****),<0.01 (**), <0.0001 (****) |
| 1A, B | Is only | 1.8 | *w;+;dHb9-GAL4/UAS-BoNT-C* | 1.16 (±0.03) | 121.13 (±7.95) | 128.28 (±4.32) | 2.22 (±0.23) | 8.31 (±0.55) | -65.12 (±1.68) | 11 | - |
| 1A, B | IIA, Is only | 1.8 | *w;GluRIIA*$^{pv3}$*;dHb9-GAL4/UAS-BoNT-C* | 0.53 (±0.01) | 71.93 (±2.52) | 136.08 (±5.45) | 1.28 (±0.19) | 7.99 (±0.65) | -67.80 (±1.43) | 8 | <0.0001 (****), <0.0001 (****), 0.25 (ns) |
| 1E | Ib only (Is> BoNT) | 0.4 | *w;+;R27E09-GAL4/UAS-BoNT-C* | 0.64 (±0.02) | 9.01 (±1.98) | 14.29 (±3.45) | 1.30 (±0.24) | 11.54 (±0.69) | -67.92 (±1.26) | 11 | - |
| 1E | IIA, Ib only (Is> BoNT) | 0.4 | *w;GluRIIA*$^{pv3}$*;R27E09-GAL4/UAS-BoNT-C* | 0.39 (±0.01) | 11.28 (±1.50) | 28.22 (±3.05) | 1.24 (±0.24) | 8.52 (±0.51) | -65.18 (±2.09) | 14 | <0.0001 (****), 0.36 (ns), <0.01 (**) |
| 1E | Ib only | 1.0 | *w;+;R27E09-GAL4/UAS-BoNT-C* | 0.67 (±0.03) | 36.97 (±1.67) | 55.62 (±2.48) | 1.88 (±0.17) | 9.08 (±0.78) | -69.28 (±2.21) | 13 | - |
| 1E | IIA, Ib only | 1.0 | *w;GluRIIA*$^{pv3}$*;R27E09-GAL4/UAS-BoNT-C* | 0.36 (±0.02) | 55.21 (±1.74) | 155.71 (±10.38) | 1.96 (±0.21) | 8.29 (±0.61) | -63.57 (±1.68) | 9 | <0.0001 (****), <0.0001 (****), <0.001 (***) |
| 1E | Ib only | 3.0 | *w;+;R27E09-GAL4/UAS-BoNT-C* | 0.70 (±0.03) | 107.24 (±10.69) | 152.18 (±16.26) | 1.86 (±0.14) | 8.81 (±0.94) | -65.24 (±2.09) | 8 | - |
| 1E | IIA, Ib only | 3.0 | *w;GluRIIA*$^{pv3}$*;R27E09-GAL4/UAS-BoNT-C* | 0.36 (±0.02) | 107.38 (±10.75) | 307.45 (±38.85) | 1.38 (±0.20) | 9.98 (±1.25) | -66.51 (±2.55) | 13 | <0.0001 (****), >0.99 (ns), <0.01 (**) |
| 1E | Ib only | 6.0 | *w;+;R27E09-GAL4/UAS-BoNT-C* | 0.71 (±0.02) | 113.34 (±6.95) | 158.47 (±9.63) | 1.98 (±0.21) | 11.24 (±1.12) | -68.79 (±2.05) | 13 | - |
| 1E | IIA, Ib only | 6.0 | *w;GluRIIA*$^{pv3}$*;R27E09-GAL4/UAS-BoNT-C* | 0.37 (±0.02) | 115.07 (±4.32) | 312.43 (±7.61) | 1.89 (±0.09) | 7.28 (±0.93) | -64.81 (±2.37) | 10 | <0.0001 (****), 0.97 (ns), <0.0001 (****) |
| 1F | Is only (Ib>BoNT) | 0.4 | *w;+;dHb9-GAL4/UAS-BoNT-C* | 1.31 (±0.06) | 25.89 (±1.64) | 20.60 (±2.77) | 1.32 (±0.19) | 10.91 (±0.75) | -67.90 (±1.53) | 8 | - |
| 1F | IIA, Is only (Ib>BoNT) | 0.4 | *w;GluRIIA*$^{pv3}$*; dHb9-GAL4/UAS-BoNT-C* | 0.60 (±0.05) | 14.17 (±1.99) | 23.59 (±2.40) | 1.20 (±0.07) | 11.21 (±0.88) | -63.48 (±0.57) | 7 | <0.0001 (****), <0.001 (***), 0.50 (ns) |
| 1F | Is only | 1.0 | *w;+;dHb9-GAL4/UAS-BoNT-C* | 1.20 (±0.04) | 88.62 (±4.72) | 74.63 (±4.72) | 1.21 (±0.12) | 6.54 (±0.42) | -67.28 (±2.41) | 10 | - |
| 1F | IIA, Is only | 1.0 | *w;GluRIIA*$^{pv3}$*;dHb9-GAL4/UAS-BoNT-C* | 0.57 (±0.02) | 47.66 (±0.49) | 85.61 (±3.77) | 1.56 (±0.08) | 8.65 (±0.57) | -63.86 (±1.72) | 11 | <0.0001 (****), <0.0001 (****), 0.28 (ns) |
| 1F | Is only | 3.0 | *w;+;dHb9-GAL4/UAS-BoNT-C* | 1.26 (±0.07) | 136.03 (±8.34) | 114.11 (±5.97) | 1.42 (±0.29) | 7.52 (±0.66) | -69.92 (±2.67) | 8 | - |
| 1F | IIA, Is only | 3.0 | *w;GluRIIA*$^{pv3}$*;dHb9-GAL4/UAS-BoNT-C* | 0.59 (±0.05) | 125.28 (±7.02) | 225.12 (±29.27) | 2.24 (±0.66) | 9.75 (±0.74) | -64.83 (±2.07) | 7 | <0.0001 (****), <0.0001 (****), <0.0001 (****) |
| 1F | Is only | 6.0 | *w;+;dHb9-GAL4/UAS-BoNT-C* | 1.27 (±0.03) | 132.46 (±7.41) | 104.89 (±6.42) | 1.38 (±0.14) | 11.25 (±0.62) | -62.69 (±2.55) | 12 | - |
| 1F | IIA, Is only | 6.0 | *w;GluRIIA*$^{pv3}$*;dHb9-GAL4/UAS-BoNT-C* | 0.59 (±0.02) | 131.81 (±1.97) | 227.99 (±7.69) | 1.71 (±0.13) | 8.00 (±0.71) | -63.54 (±2.05) | 10 | <0.0001 (****), <0.99 (ns), <0.0001 (****) |

| Figure | Label | [$Ca^{2+}$] (mM) | Genotype | PhTx | mEPSP amp(mV) | EPSC amp (-nA) | QC | mEPSP freq (Hz) | $R_{in}$ (MΩ) | $V_{rest}$ (mV) | n | One-way ANOVA, P Value: mEPSP, EPSC, QC. |
|---|---|---|---|---|---|---|---|---|---|---|---|---|
| 2A, B | wild type | 1.8 | *$w^{1118}$* | - | 1.01 (±0.03) | 180.70 (±10.79) | 171.00 (±10.83) | 4.45 (±0.27) | 8.75 (±0.41) | -64.22 (±1.01) | 8 | - |
| 2A, B | wild type+ PhTx | 1.8 | *$w^{1118}$* | + | 0.57 (±0.02) | 176.11 (±8.40) | 311.48 (±20..38) | 3.00 (±0.36) | 8.85 (±0.56) | -67.29 (±2.32) | 9 | <0.0001 (****), 0.93 (ns), <0.001 (***) |
| 2A, B | Ib only | 1.8 | *w;+;R27E09-GAL4/UAS-BoNT-C* | - | 0.59 (±0.06) | 52.35 (±4.15) | 87.60 (±5.90) | 1.98 (±0.22) | 9.78 (±0.62) | -72.86 (±1.81) | 11 | - |
| 2A, B | Ib only+ PhTx | 1.8 | *w;+;R27E09-GAL4/UAS-BoNT-C* | + | 0.35 (±0.02) | 31.41 (±2.20) | 92.68 (±11.06) | 1.44 (±0.09) | 10.20 (±0.95) | -75.22 (±2.85) | 8 | <0.0001 (****),<0.01 (**), 0.93 (ns) |
| 2A, B | Is only | 1.8 | *w;+;dHb9-GAL4/UAS-BoNT-C* | - | 1.16 (±0.03) | 121.13 (±7.95) | 128.28 (±4.32) | 2.22 (±0.23) | 8.31 (±0.55) | -65.12 (±1.68) | 11 | - |
| 2A, B | Is only + PhTx | 1.8 | *w;+;dHb9-GAL4/UAS-BoNT-C* | + | 0.46 (±0.02) | 104.67 (±7.59) | 231.23 (±19.83) | 2.20 (±0.11) | 12.05 (±1.05) | -62.96 (±1.39) | 11 | <0.0001 (****), 0.15 (ns), <0.0001 (****) |
| 2E | Ib only (Is> BoNT) | 0.4 | *w;+;R27E09-GAL4/UAS-BoNT-C* | - | 0.64 (±0.02) | 9.01 (±1.98) | 14.29 (±3.45) | 1.30 (±0.24) | 11.54 (±0.69) | -67.92 (±1.26) | 11 | - |
| 2E | Ib only (Is> BoNT) + PhTx | 0.4 | *w;+;R27E09-GAL4/UAS-BoNT-C* | + | 0.36 (±0.03) | 4.38 (±0.55) | 12.04 (±1.05) | 1.43 (±0.13) | 7.50 (±0.51) | -61.89 (±0.48) | 10 | <0.0001 (****), <0.05(*), 0.82 (ns) |
| 2E | Ib only | 1.0 | *w;+;R27E09-GAL4/UAS-BoNT-C* | - | 0.67 (±0.03) | 36.97 (±1.67) | 55.62 (±2.48) | 1.88 (±0.17) | 9.06 (±0.78) | -69.28 (±2.21) | 13 | - |
| 2E | Ib only + PhTx | 1.0 | *w;+;R27E09-GAL4/UAS-BoNT-C* | + | 0.42 (±0.03) | 20.51 (±0.47) | 49.55 (±3.38) | 1.92 (±0.07) | 7.55 (±0.85) | -64.48 (±1.54) | 9 | <0.0001 (****), <0.0001 (****), 0.68 (ns) |
| 2E | Ib only | 3.0 | *w;+;R27E09-GAL4/UAS-BoNT-C* | - | 0.70 (±0.03) | 107.24 (±10.69) | 152.18 (±16.26) | 1.86 (±0.14) | 8.81 (±0.94) | -65.24 (±2.09) | 8 | - |
| 2E | Ib only + PhTx | 3.0 | *w;+;R27E09-GAL4/UAS-BoNT-C* | + | 0.44 (±0.02) | 90.84 (±5.42) | 210.07 (±16.74) | 1.79 (±0.24) | 10.85 (±0.65) | -63.94 (±1.16) | 11 | <0.0001 (****), 0.41 (ns), 0.34 (ns) |
| 2E | Ib only | 6.0 | *w;+;R27E09-GAL4/UAS-BoNT-C* | - | 0.71 (±0.02) | 113.34 (±6.95) | 158.47 (±9.63) | 1.98 (±0.21) | 11.24 (±1.12) | -68.79 (±2.05) | 13 | - |
| 2E | Ib only + PhTx | 6.0 | *w;+;R27E09-GAL4/UAS-BoNT-C* | + | 0.44 (±0.03) | 105.81 (±7.10) | 245.31 (±18.32) | 1.90 (±0.13) | 9.95 (±0.68) | -62.97 (±1.43) | 9 | <0.0001 (****), 0.64 (ns), <0.0001 (****) |
| 2F | Is only (Ib>BoNT) | 0.4 | *w;+;dHb9-GAL4/UAS-BoNT-C* | - | 1.31 (±0.06) | 25.89 (±1.64) | 20.60 (±2.77) | 1.32 (±0.19) | 10.91 (±0.75) | -67.90 (±1.53) | 8 | - |
| 2F | Is only (Ib>BoNT) + PhTx | 0.4 | *w;+;dHb9-GAL4/UAS-BoNT-C* | + | 0.61 (±0.02) | 21.90 (±2.11) | 35.12 (±2.27) | 1.65 (±0.12) | 9.55 (±1.30) | -63.81 (±2.07) | 10 | <0.0001 (****), 0.23 (ns), <0.0001 (****) |
| 2F | Is only | 1.0 | *w;+;dHb9-GAL4/UAS-BoNT-C* | - | 1.20 (±0.04) | 88.62 (±4.72) | 74.63 (±4.72) | 1.21 (±0.12) | 6.54 (±0.42) | -67.28 (±2.41) | 10 | - |
| 2F | Is only + PhTx | 1.0 | *w;+;dHb9-GAL4/UAS-BoNT-C* | + | 0.66 (±0.08) | 67.22 (±5.31) | 110.29 (±4.72) | 2.50 (±0.11) | 7.50 (±0.81) | -66.23 (±1.98) | 8 | <0.0001 (****), <0.0001 (****), <0.001 (***) |
| 2F | Is only | 3.0 | *w;+;dHb9-GAL4/UAS-BoNT-C* | - | 1.26 (±0.07) | 136.03 (±8.34) | 114.11 (±5.97) | 1.42 (±0.29) | 7.52 (±0.66) | -69.92 (±2.67) | 9 | - |
| 2F | Is only + PhTx | 3.0 | *w;+;dHb9-GAL4/UAS-BoNT-C* | + | 0.68(±0.02) | 154.28 (±22.42) | 225.12 (±29.27) | 1.77 (±0.24) | 8.54 (±0.76) | -65.10 (±2.03) | 9 | <0.0001 (****), <0.0001 (****), <0.01 (**) |
| 2F | Is only | 6.0 | *w;+;dHb9-GAL4/UAS-BoNT-C* | - | 1.27 (±0.03) | 132.46 (±7.41) | 104.89 (±6.15) | 1.38 (±0.14) | 11.25 (±0.62) | -62.69 (±2.55) | 12 | - |
| 2F | Is only + PhTx | 6.0 | *w;+;dHb9-GAL4/UAS-BoNT-C* | + | 0.64 (±0.04) | 145.79 (±10.66) | 227.99 (±7.69) | 1.77 (±0.07) | 8.85 (±0.98) | -63.27 (±2.47) | 10 | <0.0001 (****), <0.01 (**), <0.0001 (****) |

| Figure | Label | Genotype | PhTx | BRP mean int. (% change) | RBP mean int. (% change) | Cac mean int. (% change) | n (puncta): BRP, RBP, Cac | P Value (significance): BRP density (paired T-test), mean intensity of BRP, RBP, Cac (Mann-Whitney -test) |
|---|---|---|---|---|---|---|---|---|

| | | | | | | | | |
|---|---|---|---|---|---|---|---|---|
| 3B | Ib | *Cac*$^{sfGFP-N}$;+;+ | - | - | - | - | 283, 333, 305 | - |
| 3B | IIA, Ib | *Cac*$^{sfGFP-N}$;*GluRIIA*$^{pv3}$;+ | - | 201.2 (±4.22) | 126.1 (±3.89) | 177.3 (±3.85) | 238, 204, 252 | <0.0001 (****), <0.0001 (****), <0.0001 (****) |
| 3D | Is | *Cac*$^{sfGFP-N}$;+;+ | - | - | - | - | 148, 113, 158 | - |
| 3D | Is + Phtx | *Cac*$^{sfGFP-N}$;+;+ | + | 158.6 (±6.52) | 149.1 (±3.32) | 138.4 (±3.77) | 140,134,177 | <0.0001 (****), <0.0001 (****), <0.0001 (****) |

| Figure | Label | Genotype | PhTx | Brp nanomod./Ring | n (puncta) | P Value (significance): Mann-Whitney test |
|---|---|---|---|---|---|---|
| 4B | Ib | *Cac*$^{sfGFP-N}$;+;+ | - | 5.25 (±0.20) | 21 | - |
| 4B | IIA, Ib | *Cac*$^{sfGFP-N}$;*GluRIIA*$^{pv3}$;+ | - | 7.01 (±0.35) | 12 | 0.0002 (***) |
| 4H | Is | *Cac*$^{sfGFP-N}$;+;+ | - | 4.56 (±0.21) | 20 | - |
| 4H | Is+ PhTx | *Cac*$^{sfGFP-N}$;+;+ | + | 3.99 (±0.38) | 8 | 0.2525 (ns) |

| Figure | Label | Genotype | PhTx | Cac area (nm$^2$) | BRP ring area (nm$^2$) | Cac/BRP area ratio (%) | n (puncta): BRP, Cac | Unpaired T-test ,P Value (significance): BRP ring area (μm$^2$), RBP/BRP area ratio (%), nanomodules/ring, |
|---|---|---|---|---|---|---|---|---|
| 4C, D | Ib | *Cac*$^{sfGFP-N}$;+;+ | - | 23392 (±1267) | 64359 (±2507) | 36.11 (±1.317%) | 43, 48 | - |
| 4C, D | IIA, Ib | *Cac*$^{sfGFP-N}$;*GluRIIA*$^{pv3}$;+ | - | 29872 (±1665) | 78405 (±3469) | 39.20 (±1.635%) | 47, 52 | 0.2821 (ns) |
| 4I, J | Is | *Cac*$^{sfGFP-N}$;+;+ | - | 23712 (±1493) | 44762 (±3269) | 55.52 (±2.485%) | 19, 21 | - |
| 4I, J | Is + PhTx | *Cac*$^{sfGFP-N}$;+;+ | + | 17877 (±935.2) | 46429 (±2864) | 39.68 (±2.526%) | 17, 19 | 0.0003 (***) |

| Figure | Label | Genotype | PhTx | RBP area (nm$^2$) | BRP ring area (nm$^2$) | RBP/BRP area ratio (%) | n (puncta): BRP, RBP | Unpaired T-test ,P Value (significance): BRP ring area (μm$^2$), RBP/BRP area ratio (%), nanomodules/ring, |
|---|---|---|---|---|---|---|---|---|
| 4C, D | Ib | *Cac*$^{sfGFP-N}$;+;+ | - | 36219 (±1668) | 62727 (±3564) | 60.28 (±2.015%) | 25, 25 | - |
| 4C, D | IIA, Ib | *Cac*$^{sfGFP-N}$;*GluRIIA*$^{pv3}$;+ | - | 48696 (±2290) | 79621 (±3259) | 62.26 (±1.365%) | 42, 42 | 0.6785 (ns) |
| 4I, J | Is | *Cac*$^{sfGFP-N}$;+;+ | - | 34545 (±2081) | 55028 (±2738) | 62.27 (±1.888%) | 20, 20 | - |
| 4I, J | Is + PhTx | *Cac*$^{sfGFP-N}$;+;+ | + | 30459 (±1568) | 50574 (±2468) | 62.42 (±1.719%) | 24, 24 | 0.9932 (ns) |

| Figure | Label | [Ca$^{2+}$] (mM) | Genotype | PhTx | Indicator | dR/R | Decay (ms) | Rise (ms) | n | Paired T-test, P Value (significance): ratio of indicators |
|---|---|---|---|---|---|---|---|---|---|---|
| 5B, C | Ib | 1.8 | *w;+;OK319-GAL4/UAS-Syt::mScarlet::GCaMP8f* | - | GCaMP8f /mScarlet | 0.34 (±0.01) | 75.4 (±3.1) | 34.7 (±1.4) | 17 | - |
| 5B, C | IIA, Ib | 1.8 | *w; +; OK319-GAL4,GluRIIA*$^{pv3}$*/UAS-Syt::mScarlet::GCaMP8f* | - | GCaMP8f /mScarlet | 0.51 (±0.03) | 69.2 (±2.0) | 28.6 (±0.8) | 25 | <0.0001 (****) |
| 5E, F | Is | 1.8 | *w;+;OK319-GAL4/UAS-Syt::mScarlet::GCaMP8f* | - | GCaMP8f /mScarlet | 0.76 (±0.05) | 78.8 (±3.5) | 34.7 (±1.1) | 19 | - |
| 5E, F | Is + PhTx | 1.8 | *w;+;OK319-GAL4/UAS-Syt::mScarlet::GCaMP8f* | + | GCaMP8f /mScarlet | 0.79 (±0.03) | 81.1 (±2.3) | 32.0 (±0.9) | 24 | 0.909 (ns) |

| Figure | Label | [Ca$^{2+}$] (mM) | Genotype | PhTx | Cum. EPSC(-nA) | Est. RRP size | $R_{in}$ (MΩ) | $V_{rest}$ (mV) | n | Unpaired T-test, P Value (significance): Cum. EPSC, RRP size |
|---|---|---|---|---|---|---|---|---|---|---|
| 6A, B | Ib only | 3.0 | *w;+;R27E09-GAL4/UAS-BoNT-C* | - | 586.6 (±47.72) | 952.3 (±77.46) | 6.31 (±0.67) | -66.78 (±1.55) | 15 | - |
| 6A, B | IIA, Ib | 3.0 | *w;GluRIIA*$^{pv3}$*;R27E09-GAL4/UAS-BoNT-C* | - | 351.4 (±33.22) | 994.3(±94.01) | 5.81 (±0.38) | -63.34 (±2.71) | 12 | <0.001 (***), 0.73 (ns) |
| 6C, D | Is only | 3.0 | *w;+;dHb9-GAL4/UAS-BoNT-C* | - | 688.5 (±70.56) | 719.7 (±73.75) | 8.20 (±0.56) | -65.99 (±1.80) | 10 | - |
| 6C, D | Is + PhTx | 3.0 | *w;+;dHb9-GAL4/UAS-BoNT-C* | + | 568.3 (±85.00) | 1566.00 (±160.4) | 7.25 (±0.77) | -67.83 (±1.93) | 9 | 0.28 (ns),<0.001 (***) |

| Figure | Label | Genotype | PhTx | Release site # | $R_{in}$ (MΩ) | $V_{rest}$ (mV) | n | Unpaired T-test, P Value (significance): Release site # |
|---|---|---|---|---|---|---|---|---|
| 7A, B, C | Ib | *w;+;R27E09-GAL4/UAS-BoNT-C* | - | 518.8 (±47.72) | 6.00 (±0.5) | -67.8 (±1.02) | 9 | - |
| 7A, B, C | IIA, Ib | *w;GluRIIA$^{pv3}$;R27E09-GAL4/UAS-BoNT-C* | - | 794.4 (±60.76) | 5.6 (±0.55) | -66 (±0.83) | 8 | 0.0026 (**) |
| 7G, H, I | Is | *w;+;dHb9-GAL4/UAS-BoNT-C* | - | 329.1 (±31.27) | 7.5 (±0.89) | -66.9(±1.14) | 10 | - |
| 7G, H, I | Is + PhTx | *w;+;dHb9-GAL4/UAS-BoNT-C* | + | 533.3 (±61.83) | 5.5 (±0.37) | -67.7(±1.47) | 7 | 0.0057 (**) |

| Figure | Label | Genotype | PhTx | Unc13A mean int. (% change) | Unc13A area (μm$^2$) | n (puncta) | P Value (significance): mean intensity of Unc13A (Mann-Whitney test), Unc13A area (Mann-Whitney test), |
|---|---|---|---|---|---|---|---|
| 7E, F | Ib | *Cac$^{sfGFP-N}$;+;+* | - | - | 0.073 (±0.003) | 289, 151 | - |
| 7E, F | IIA, Ib | *Cac$^{sfGFP-N}$;GluRIIA$^{pv3}$;+* | - | 151.4 (±2.10) | 0.086 (±0.003) | 232, 158 | <0.0001 (****); 0.0043 (**) |
| 7K, L | Is | *Cac$^{sfGFP-N}$;+;+* | - | - | 0.039 (±0.003) | 123, 49 | - |
| 7K, L | Is + Phtx | *Cac$^{sfGFP-N}$;+;+* | + | 122.6 (±4.82) | 0.044 (±0.003) | 124, 61 | 0.0039 (**); 0.3289 (ns) |

| Figure | Label | [$Ca^{2+}$] (mM) | Genotype | PhTx | EGTA-AM | EPSC (nA) | % of baseline (-EGTA) | $R_{in}$ (MΩ) | $V_{rest}$ (mV) | n | Unpaired T-test, P Value (significance): % baseline (-EGTA) |
|---|---|---|---|---|---|---|---|---|---|---|---|
| 8A, B | Ib only | 1.8 | *w;+;R27E09-GAL4/UAS-BoNT-C* | - | - | -71.164(±4.06) | - | 7.27 (±0.51) | -69.51 (±1.02) | 17 | - |
| 8A, B | Ib only | 1.8 | *w;+;R27E09-GAL4/UAS-BoNT-C* | - | 50 μM | -43.41 (±2.72) | 61.00 (±4.10) | 6.44 (±0.57) | -67.51 (±2.61) | 8 | - |
| 8A, B | IIA, Ib | 1.8 | *w;GluRIIA$^{pv3}$;R27E09-GAL4/UAS-BoNT-C* | - | - | -84.82 (±4.02) | - | 7.52 (±0.68) | -63.42 (±1.57) | 8 | - |
| 8A, B | IIA, Ib | 1.8 | *w;GluRIIA$^{pv3}$;R27E09-GAL4/UAS-BoNT-C* | - | 50 μM | -80.98 (±5.65) | 79.71 (±3.975) | 6.51 (±0.69) | -65.88 (±1.24) | 14 | <0.001 (**) |
| 8C, D | Is only | 1.8 | *w;+;dHb9-GAL4/UAS-BoNT-C* | - | - | -145.80 (±6.13) | - | 6.74 (±0.81) | -64.11 (±1.71) | 8 | - |
| 8C, D | Is only | 1.8 | *w;+;dHb9-GAL4/UAS-BoNT-C* | - | 50 μM | -108.05 (±5.28) | 76.11 (±3.926) | 7.78 (±0.95) | -64.27 (±2.84) | 10 | - |
| 8C, D | Is + PhTx | 1.8 | *w;+;dHb9-GAL4/UAS-BoNT-C* | + | - | -152.80(±8.50) | - | 9.24 (±0.88) | -67.85(±1.79) | 8 | - |
| 8C, D | Is + PhTx | 1.8 | *w;+;dHb9-GAL4/UAS-BoNT-C* | + | 50 μM | -88.55 (±3.90) | 58.38 (±2.962) | 7.76 (±0.61) | -68.82 (±1.82) | 10 | <0.001 (**) |

| Figure | Label | Genotype | +PhTx | merged BRP rings | total BRP rings | n (boutons) | Unpaired T-test, P Value (significance): Brp puncta #/M6 NMJ |
|---|---|---|---|---|---|---|---|
| S1B | Ib | *w$^{1118}$* | - | 3.947 (±0.4152) | 18.89 (±1.139) | 25 | - |
| S1B | IIA, Ib | *GluRIIA$^{pv3}$* | - | 5.267 (±0.4306) | 18 (±1.480) | 15 | <0.0001 (****) |
| S1D | Is | *w$^{1118}$* | - | 1.156 (±0.1687) | 6.750 (±0.5803) | 39 | - |
| S1D | Is + PhTx | *w$^{1118}$* | + | 1.650 (±0.2325) | 7.650 (±0.5771) | 27 | 0.1188 (ns) |
