## Supplementary material for "Distinct input-specific mechanisms enable presynaptic homeostatic plasticity": Key Resources Table

| REAGENT/RESOURCE | | SOURCE | IDENTIFIER |
|---|---|---|---|
| **Chemicals, peptides, and recombinant proteins** | | | |
| Philanthotoxin-433 | | Sigma-Aldrich | 276684-27-6 |
| EGTA-AM | | Sigma-Aldrich | 99590-86-0 |
| **Antibodies** | **Dilution** | | |
| Mouse anti-BRP (nc82) | 1:250 | Developmental Studies Hybridoma Bank (DSHB) | AB_2314866 |
| Guinea pig anti-RBP | 1:100 | (He et al., 2023) | |
| Guinea pig anti-Unc13A | 1:100 | (Böhme et al., 2016) | |
| Chicken anti-GFP | 1:400 | Aves Labs | GFP-1020 |
| Alexa Fluor 488 conjugated Goat anti-Horseradish Peroxidase | 1:400 | Jackson ImmunoReserach Laboratories (Jackson) | 123-545-021 |
| Alexa Fluor 488 conjugated secondary antibodies | 1:400 | Jackson | 715-545-150 |
| Cy3-conjugated secondary antibodies | 1:400 | Jackson | 106-165-003 |
| Alexa Fluor 594 conjugated secondary antibodies | 1:400 | Jackson | 106-585-003<br>103-585-155 |
| Alexa Fluor 647 conjugated Goat anti-Horseradish Peroxidase | 1:400 | Jackson | 123-605-021 |
| STAR RED conjugated secondary antibodies | 1:200 | Abberior | STRED-1001<br>STRED-1006 |
| **Drosophila Strains** | | | |
| UAS-Syt::mScarlet::GCaMP8f | | (Li et al., 2021) | |
| UAS-BoNT-C | | (Han et al., 2022) | |
| *Cac*$^{sfGFP\text{-}N}$ | | (Gratz et al., 2019) | |
| OK319-Gal4 | | (Beck et al., 2012) | |
| dHb9-GAL4 (Ib-Gal4) | | BDSC | 83004 |
| GMR27E09-GAL4 (Is-Gal4) | | BDSC | 49227 |
| *GluRIIA*$^{pv3}$ | | (Han et al., 2023) | |
| *w*$^{1118}$ | | BDSC | 5905 |
| **Software and Algorithms** | | | |
| NIS Elements software | | Nikon | 4.51.01 |
| Huygens Essential | | SVI | 22.10 |
| Axon pCLAMP | | Molecular Devices | 10.7 |
| MiniAnalysis | | Synaptosoft | 6.0.3 |
| GraphPad Prism | | GraphPad | 8.0.1 |
